# Allelic imbalance reveals widespread germline-somatic regulatory differences and prioritizes risk loci in Renal Cell Carcinoma

**DOI:** 10.1101/631150

**Authors:** Alexander Gusev, Sandor Spisak, Andre P. Fay, Hallie Carol, Kevin C Vavra, Sabina Signoretti, Viktoria Tisza, Mark Pomerantz, Forough Abbasi, Ji-Heui Seo, Toni K. Choueiri, Kate Lawrenson, Matthew L Freedman

## Abstract

Determining the function of non-coding regulatory variants in cancer is a key challenge transcriptional biology. We investigated genetic (germline and somatic) determinants of regulatory mechanisms in renal cell carcinoma (RCC) using H3K27ac ChIP-seq data in 10 matched tumor/normal samples and RNA-seq data from 496/66 tumor/normal samples from The Cancer Genome Atlas (TCGA). Unsupervised clustering of H3K27ac activity cleanly separated tumor from normal individuals, highlighting extensive epigenetic reprogramming during transformation. We developed a novel method to test each chromatin feature for evidence of an allele-specific quantitative trait locus (asQTL) and evaluate tumor/normal differences in allele-specificity (d-asQTLs) while modelling local structural variation and read overdispersion. At an FDR of 5%, we identified 1,356 unique asQTL chromatin peaks in normal tissues; 2,868 in tumors; and 1,054 d-asQTLs (primarily imbalanced in tumor). The d-asQTL peaks were significantly enriched for RCC genome-wide association study (GWAS) heritability (32x, P=1.8×10^−3^), more so than any other functional feature including all H3K27ac peaks (12x), super-enhancers (5x), and asQTL genes (4x). Intersection of asQTLs with RCC GWAS loci identified putative functional features for 6/17 known loci including tumor-specific activity at SCARB1, a cholesterol metabolism mediator, which has recently been implicated in RCC progression. We validated the asQTL variant through CRISPR interference (CRISPRi) and demonstrated a concomitant allelic effect on the overlapping enhancer and on downstream SCARB1 expression. Knockdowns of master transcription factors (TFs) involved in the hypoxia pathway altered the expression of SCARB1 in a kidney cancer cell line, consistent with a variant-TF interaction. Genome-wide, d-asQTLs were significantly enriched for tumor-specific binding of hypoxic transcription factors, implicating a more general mechanism for polygenic germline-somatic interaction.

## INTRODUCTION

Genome-wide association studies (GWASs) of cancer have been highly successful at identifying hundreds of novel risk loci, but elucidating the underlying mechanisms has posed a significant challenge. As with many complex traits, cancer risk is driven by thousands of variants of individually small effect, the majority of which reside in non-coding regions of the genome ^1–3^. While coding variation is largely interpretable, the rules governing the functional impact of non-coding, regulatory variants are poorly understood.

Identification of expression quantitative trait loci (eQTLs) in populations has proven to be an effective way to connect non-coding variants to their putative susceptibility genes. Recent analyses of population-based epigenomes have further revealed that eQTL SNPs are often also associated with variation in nearby epigenomic features (such as active enhancers marked by H3K27ac) in coordinated regulatory modules ^4–9^. Together with the observation that GWAS heritability is enriched at variants that lie in tissue-relevant epigenomic features ^7, 10–13^, a coherent model is emerging wherein GWAS variants typically alter transcription factor binding which modulates the local chromatin landscape and the expression of a gene or nearby susceptibility gene (or genes), leading to changes in disease risk. Discovery of specific genetic variants that putatively alter gene expression and epigenomic regulation can thus prioritize targets for experimental follow-up and therapeutics.

A major barrier to integrating such regulatory SNPs with GWAS data is the need for a large number of samples with functional measurements, which is further exacerbated by confounding from technical and environmental noise ^14^. High-throughput sequencing platforms offer an opportunity to mitigate these issues by leveraging within-sample variation through allele-specificity of ChIP-/RNA-seq reads. A heterozygous variant within a given functional feature (enhancer, exon, etc.) can serve as a reporter of *cis*-regulatory changes, where the number of reads mapping to each allele is an estimate of the relative activity at the respective haplotype ^15–18^. ChIP-seq allele-specific activity in multiple individuals is reflective of an underlying *cis*-regulatory variant in the population, empowering discovery of quantitative trait loci (QTLs). Additionally, consistent allele-specific activity for the same allele in multiple individuals is evidence that the variant drives (or tags) the underlying regulatory change, thereby enhancing the fine-mapping of causal variants. Allele-specific tests have several appealing properties relative to traditional association-based QTL studies. First, modelling the distribution of individual reads rather than a single quantity for total activity yields a statistical test where power tracks with read coverage rather than sample size, enabling robust QTL discovery from very small studies. Second, the allele-specific test is less susceptible to technical and environmental artifacts by leveraging *within*-sample differences in allelic activity rather than *between*-sample differences in overall activity (which reflect both *cis*-, *trans*-, and technical variance). This property is particularly important when identifying differential allele-specific quantitative trait loci between tumor and normal states, where large-scale genetic and epigenetic changes in the global regulatory landscape in the tumor make identification and correction of artifacts is challenging. The traditional approach of inferring hidden confounders using principal component or factor analysis ^19^ typically separates all tumor/normal samples and would thus result in an undefined or unstable interaction test. The allelic imbalance model has been applied extensively in studies of normal expression, but no formal interaction test has been proposed, and the previous applications to cancer have not modelled tumor-specific structural variation that can introduce biases ^20–22^.

## RESULTS

### The landscape of chromatin activity in RCC

We performed chromatin immunoprecipitation followed by high-throughput sequencing (ChIP-seq) assay of the H3K27ac histone modification (marking active enhancers and promoters) in 10 matched RCC tumor and normal samples. The same set of individuals had also been analyzed by The Cancer Genome Atlas (TCGA) with assays for germline genotypes, copy number alterations, RNA-seq, and array-based methylation. A total of 131,815 broad chromatin peaks were identified across all samples using the MACS algorithm (61,706 in normal tissues; 72,273 in tumors) of which 102,375 were active in at least two individuals. Unsupervised clustering of correlation in activity across peaks completely separated tumor/normal samples (**Figure 1a**). We identified 21,575 chromatin peaks that were significantly differentially active between tumor/normal at 1% FDR (9,491 more active in tumor and 12,084 more active in normal). As expected, tumor-specific peaks were correlated with significantly higher expression of nearby genes, and significantly lower methylation of overlapping probes in the full TCGA cohort (**Figure 1c,d**). We additionally detected 3,648 distal super-enhancers, defined as significant clusters of H3K27ac peaks not overlapping promoters using established detection criteria ^23^. Six super-enhancers were somatically acquired in all tumors and unobserved in all normal samples (**Figure 1b**) and 757 (824) were significantly more active in tumor (normal) at 5% FDR (see Methods). The strongest tumor-specific super-enhancer overlapped *EGLN3*, which is the most upregulated gene in the hypoxia pathway in RCC ^24^. The strongest normal-specific super-enhancer overlapped *TMPRSS2*, which is involved in a key fusion biomarker in prostate cancer ^25^ but has not been previously connected to RCC. Notably, the most significant super-enhancers observed in tumors were typically not tumor-specific, underscoring the utility of comparing with matched normal samples.

**Figure 1.**
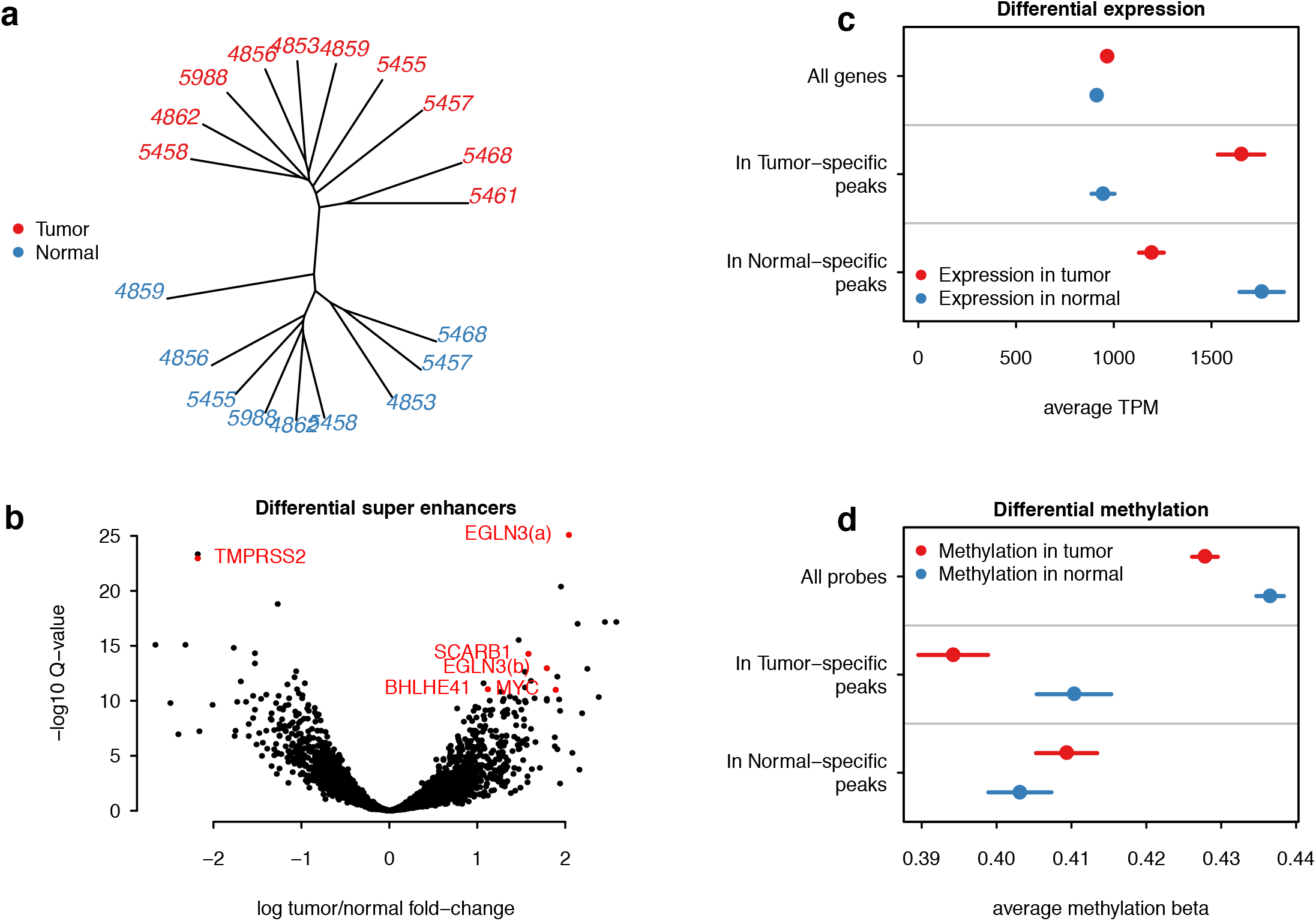
Landscape of chromatin activity in RCC. (**a**) Unsupervised clustering of H3k27ac peak activity in tumor/normal samples. Numbers indicate TCGA sample identifiers, red=tumor, blue=normal. (**b**) Volcano plot of differential super enhancer activity. Super-enhancers overlapping known oncogenes are highlighted in red (e.g. TMPRSS2 lost in all tumors; EGLN3 gained in all tumors). (**c**) Tumor/normal expression (x-axis: in transcripts per million, TPM) of genes downstream to tumor-/normal-specific H3k27ac peaks. Mean (point) and standard error (bar) shown for expression in tumor (red) and normal (blue). Tumor-specific peaks are near genes overexpressed in tumor. (**d**) Tumor/normal array-based methylation effect-size (normalized to 0-1) of probes overlapping tumor-/normal-specific H3k27ac peaks. Mean (point) and standard error (bar) shown for methylation in tumor (red) and normal (blue). Tumor-specific peaks overlap probes that are significantly hypo-methylated in tumor.

### Overview of allelic imbalance methods

To identify *cis* regulatory variants in tumor/normal samples and identify differences between these contexts we developed the *stratified Allele-Specific* (*stratAS*) test (see Methods). Allele-specific analyses pose unique challenges, particularly their vulnerability to false-positive associations when the read distribution is not modelled properly. Multiple studies have shown that molecular sequence reads have greater variability than expected from a binomial distribution ^15–17^. In tumor data, we also found this variability to be strongly dependent on local somatic copy number alterations (SCNA) (**Supplementary Table S1**). We account for these confounders by modelling the read distribution as a beta-binomial, with over-dispersion parameters inferred for each sample and each local CNA change (**Figure 2a**). *stratAS*, like other AS approaches ^15, 17^, incorporates haplotype phase information to jointly test all heterozygous variants within a functional feature (peak or multiple exons). This can increase power substantially for analysis of broad H3K27ac peaks which contained 8.3 SNPs on average in our data. Lastly, *stratAS* identifies tumor/normal differential regulatory variants by evaluating a likelihood ratio test for difference in the underlying allelic fraction (**Figure 2b**). This represents an important difference from previous strategies that independently test imbalance in each condition and then look for features that are marginally significant in only one condition ^21^. For features where activity (and thus coverage) systematically changes between conditions, the marginal significance-based approach may identify false differences simply due to greater detection power in one condition. In contrast, the *stratAS* test for a statistical difference in the allelic fraction accounts for read distributions in both conditions.

**Figure 2.**
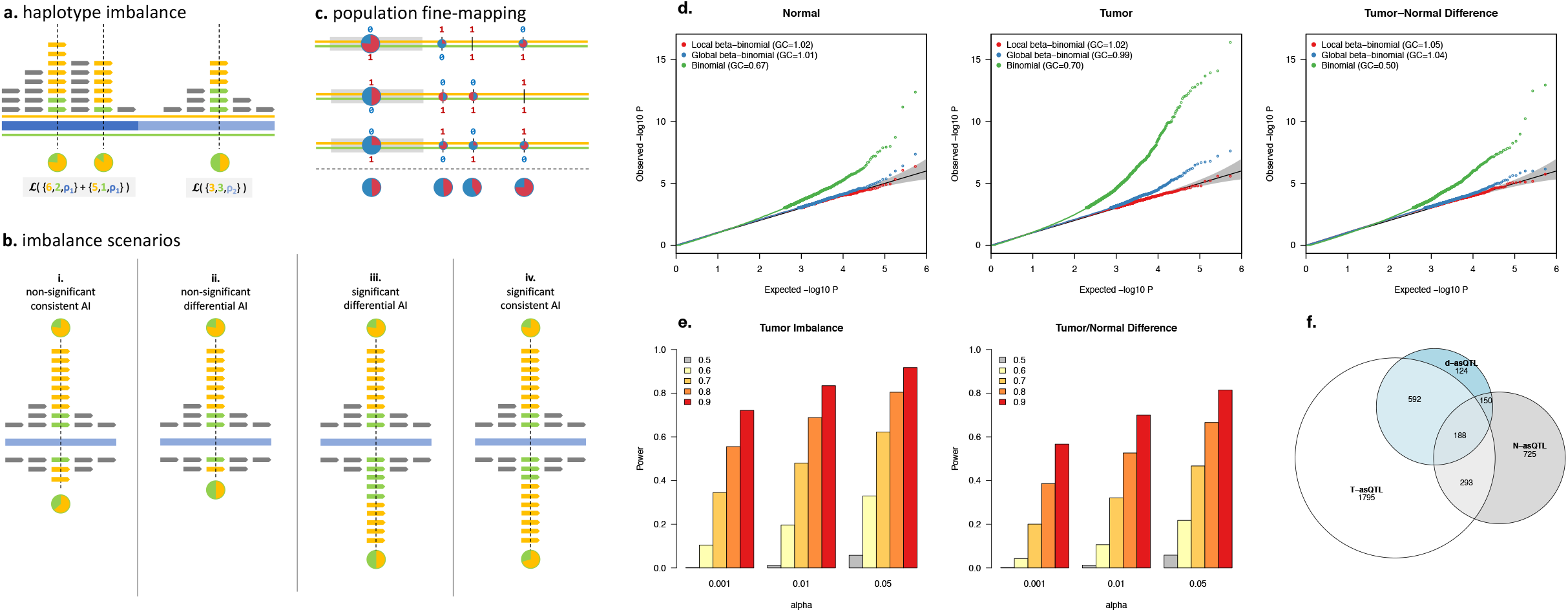
Allelic imbalance model. (**a**) Haplotype imbalance in a single individual. Orange/green indicate two (maternal/paternal) haplotypes; reads mapping to those haplotypes at heterozygous sites (dashed vertical lines); and allelic fraction (pie chart). Blue shaded bars indicate local CNA boundaries and corresponding over-dispersion parameter ρ_*,*ij*_. Likelihood computation across the haplotype shown in gray blocks. (**b**) Four possible imbalance scenarios in a single individual with two conditions (e.g. reads from tumor above the blue line, reads from normal below): (i) consistent allelic imbalance but not significant in normal due to few reads; (ii) differential allelic imbalance but not significant in normal due to few reads; (iii) differential allelic imbalance with sufficient reads to detect significant difference (d-asQTL); (iv) significant allelic imbalance that is consistent between tumor/normal. (**c**) Fine-mapping the causal regulatory variant in samples from a population. Each orange/green line pair represents a maternal/paternal haplotype for an individual. Red/blue numbers represent bi-allelic SNPs, with each pie-chart showing the allelic fraction at each individual and site (with average below the dashed line). Gray block is the exon containing the read-carrying variant, whereas the right-most SNP is the true causal variant. Haplotype-based test across individuals resolves the right-most variant as the most imbalanced. (**d**) QQ-plots for allelic imbalance test under the null of no imbalance shown for normal (left), tumor (middle), and differential (right). Traditional binomial test (green) is inflated in all scenarios; beta-binomial test with a single global over-dispersion parameter (blue) is inflated in tumor; stratAS test with CNA-specific over-dispersion parameter is well-calibrated in all scenarios. (**e**) Power to detect allelic imbalance using the stratAS test. Y-axis is fraction of loci detected at a given significance threshold (x-axis) and true imbalance (bar color; gray = no imbalance, yellow-red = increasing imbalance). Left panel shows simulation in tumor, right panel shows tumor/normal difference test (where imbalance in normal is always null). (**f**) Number and overlap of significant asQTLs (at 5% FDR) detected in each state.

We evaluated the performance of *stratAS* in simulations using the actual haplotypes, ChIP-seq read distributions, and chromatin peaks observed in the 10 H3K27ac samples. We compared *stratAS* to a beta-binomial model using a single global overdispersion parameter (equivalent to the WASP approach ^17^), as well as a standard binomial model which has been previously recommended for analysis of RNA-seq ^26^ and for which tumor/normal differences can be evaluated by a *χ*^2^ test. In normal samples, both *stratAS* and the global beta-binomial tests were well-controlled under the null, with little confounding from structural variation (**Figure 2d**). However, in tumor samples, only the *stratAS* model sufficiently addressed confounding due to SCNAs, whereas the global model was slightly inflated at high coverage. In contrast, the standard binomial test poorly modelled the data across all states and exhibited both a deflation for low counts (median *λ_GC_* 0.5-0.7) and inflation for high counts, as well as the corresponding *χ*^2^ test for tumor/normal difference. This inflation was particularly severe for tumor data, with p-values as low as 10^−15^ for an expectation of 10^−5^.

As *stratAS* was the only well-calibrated statistic, we next focused on evaluating its power to detect true regulatory effects by simulating a range of allele-specificity in the tumor data (see Methods). Overall, *stratAS* had >50% power to identify allelic fractions >0.7 in the tumor, and tumor/normal differences of >0.8 vs 0.5 at a Type I error of 0.05 (**Figure 2e**). We computed power for a simulation with 5-fold greater coverage and observed >50% power to detect allelic fractions as low as 0.6 (**Supplementary Figure S1**). Under the simplifying assumption that the true allelic fraction is the same in all individuals, this also corresponds to the expected power from a 5-fold larger study. Lastly, we confirmed that *stratAS* outperformed previously proposed interaction methods that do not model the read distribution, including testing for correlation between allele-fraction and condition (which does not model individual reads) as well as using approximate standard errors of the independent allelic fractions (**Supplementary Figure S2, Supplementary Note**).

### stratAS identifies thousands of allele-specific chromatin QTLs

We applied the *stratAS* test across the ten tumor/normal H3K27ac samples and identified thousands of variants with significant allelic imbalance. *stratAS* was applied to all heterozygous carriers within a peak that had at least six reads and at least one read for both alleles, with 54,025/131,815 meeting these criteria. For each peak, a test for imbalance was separately performed in all normal samples and all tumor samples, as well as a for difference in imbalance between the tumor/normal samples. We refer to peaks that are imbalanced in each of these states as N-asQTL, T-asQTL, and d-asQTL respectively (without distinguishing which heterozygous variant in the peak is most imbalanced). At an FDR of 5% we identified 2,868 T-asQTLs; 1,356 N-asQTLs; and 1,054 d-asQTLs (**Table 1, Figure 2f**). The largest subcategory of d-asQTLs were 592/1,054 T-asQTLs that were not N-asQTLs, in contrast to only 150/1,054 N-asQTLs that were not T-asQTLs; with the remainder either asQTLs in both states but significantly more imbalanced in one state or asQTLs non-significantly imbalanced in both states in opposite directions, producing a significant difference. To account for differences in detection power between conditions, we estimated the fraction of asQTLs discovered in one state that were nominally significant in the other state using a bootstrapped FDR approach, finding that 87% of N-asQTLs were nominal T-asQTLs, while only 53% of T-asQTLs were nominal N-asQTLs (**Supplementary Figure S3**, Methods). These state differences reflect regulatory changes that occur during neoplastic transformation, either driven by interactions with tumor-specific regulatory activity or the presence of allele-specific SCNAs that induce imbalance in chromatin activity (with *stratAS* appropriately modelling the local change in read overdispersion).

To further characterize the regulatory changes in the tumor, we investigated the relationship between T-/d- asQTLs and local features within each individual. We re-ran *stratAS* on each H3K27ac sample independently, restricted to peaks with high power to detect imbalance (having a total >50 reads across all heterozygous sites in the peak). We then built a logistic regression across all individual-peak pairs, with the presence/absence of a significant T-/d-asQTL as the outcome and various local features as predictors (mirroring similar analyses of allele-specific expression in previous work ^22^). We found that the presence of an SCNA loss was the feature most predictive of a T-asQTL (OR of 3.5, explaining 9.6% of variance) followed by the presence of an N-asQTL in the matched normal (OR of 5.24, but explaining 2.9% of variance) (**Supplementary Table S2**). SCNA loss events were even more predictive of d-asQTLs (OR=10.6), explaining 12.8% of the variance in d-asQTL events (**Supplementary Table S2**). Interestingly, SCNA gains were not associated with d-asQTLs in a joint model with all local features and explained <1% of the variance. The outsized contribution from loss events was not explained by the magnitude of events, as CN loss/gains were comparable in frequency and absolute copy number change at the tested peaks. In total, all local features explained 14-16% of the variance in T-asQTLs and d-asQTLs, suggesting that most of the imbalance either arises from local genetic polymorphisms or unmodeled confounders.

This relationship between SCNAs and asQTLs was also apparent in the population-based analysis across all individuals. Under the null where CNA and allelic imbalance is independent, the number of asQTLs detected in a locus is expected to depend only on the total allelic read coverage. In contrast, we observed a clear relationship between SCNA variability and the number of T-/d- asQTLs (**Supplementary Figure S4**). For example, the 20% of sites in the most copy number variable regions (i.e. sites with greatest absolute CN change from diploid) accounted for 50% of T-asQTLs and 60% of d-asQTLs, after matching on total coverage. In contrast, no relationship between SCNA variability and number of detected QTLs was observed in the normal samples, as expected under the null (suggesting that normal CNA polymorphisms are not noticeably contributing to imbalance in the normal state).

### Coordinated asQTL effects on chromatin activity and gene expression

We next applied *stratAS* to RNA-seq from 524 tumor samples and 66 matched normals collected by TCGA, finding a striking 5,899 out of 9,475 tested genes to exhibit allele-specific expression at 5% FDR (**Table 1**). The substantial fraction of genes under significant genetic control is consistent with recent estimates from large eQTL studies. The increased sample size allowed us to leverage haplotype information to test all polymorphisms +/-100kb of the TSS rather than just exonic read carrying markers (**Figure 2c**). Similar to the trend for allele-specific binding, 5,731 genes were imbalanced in the tumors; 2,060 imbalanced in the normals (of which 1,893 were also significantly imbalanced in tumor); and 838 with significant tumor/normal differences. Down-sampling analyses showed that hundreds of imbalanced genes could be detected at sample sizes as low as N=10, and *stratAS* consistently yielded as many significant genes as an eQTL study from 2-10x more samples, particularly in the tumor cells (**Supplementary Figure S5**). Both expression and chromatin asQTLs were enriched for expression QTL sites detected by the GTEx Consortium 14 compared to non-significant asQTLs peaks (see Methods), serving as independent replication of their regulatory activity in normal tissues (**Supplementary Figure S6**).

We evaluated the persistence of asQTLs across assays and states in five samples with both ChIP-seq and RNA-seq available. From RNA-seq, we found that 95% of N-asQTLs were nominally significant in tumor and 70% of T-asQTLs were nominally imbalanced in normal (again using a bootstrapped FDR approach; **Supplementary Figure S4**, Methods). This reflects substantially less tumor/normal variability in the cis-regulation of gene expression than of chromatin activity, as has been previously observed at the level of overall activity ^27^. Regulatory effects were also largely consistent between H3K27ac and RNA-seq, with 30-50% asQTLs identified in one assay estimated to be nominally imbalanced in the other (**Supplementary Figure S7, Supplementary Table S3**).

### asQTLs identify specific GWAS mechanisms

The highly significant enrichment for polygenic heritability for imbalanced chromatin peaks suggests that causal germline variants are likely to reside in and be mediated by imbalanced variants. We therefore investigated the presence of allelic imbalance at 17 established, genome-wide significant RCC associations. Of the 17 variants, 3/17 overlapped significant chromatin asQTLs and 5/17 overlapped significant gene asQTLs (at 5% genomewide FDR), for a total of 6/17 asQTL GWAS SNPs (**Figure 3a**). 5/17 SNPs also resided in super-enhancers, of which 4/5 were significantly differently active between tumor and normal (**Supplementary Table S4**). In total, 10/17 RCC GWAS variants could be connected to at least one functional feature, which is striking given that 6 variants were not heterozygous in the functional data and therefore could not be tested for imbalance (**Figure 3a**). In contrast, previous eQTL analysis across the same set of TCGA samples implicated only a single gene (*DPF3*) with shared eQTL effect ^1^, which was also identified through allelic imbalance in our study. The newly characterized loci highlight the increased power of our allelic imbalance approach, and sufficient sensitivity to detect disease-relevant asQTLs in just 10 tumor-normal ChIP-seq samples.

**Figure 3.**
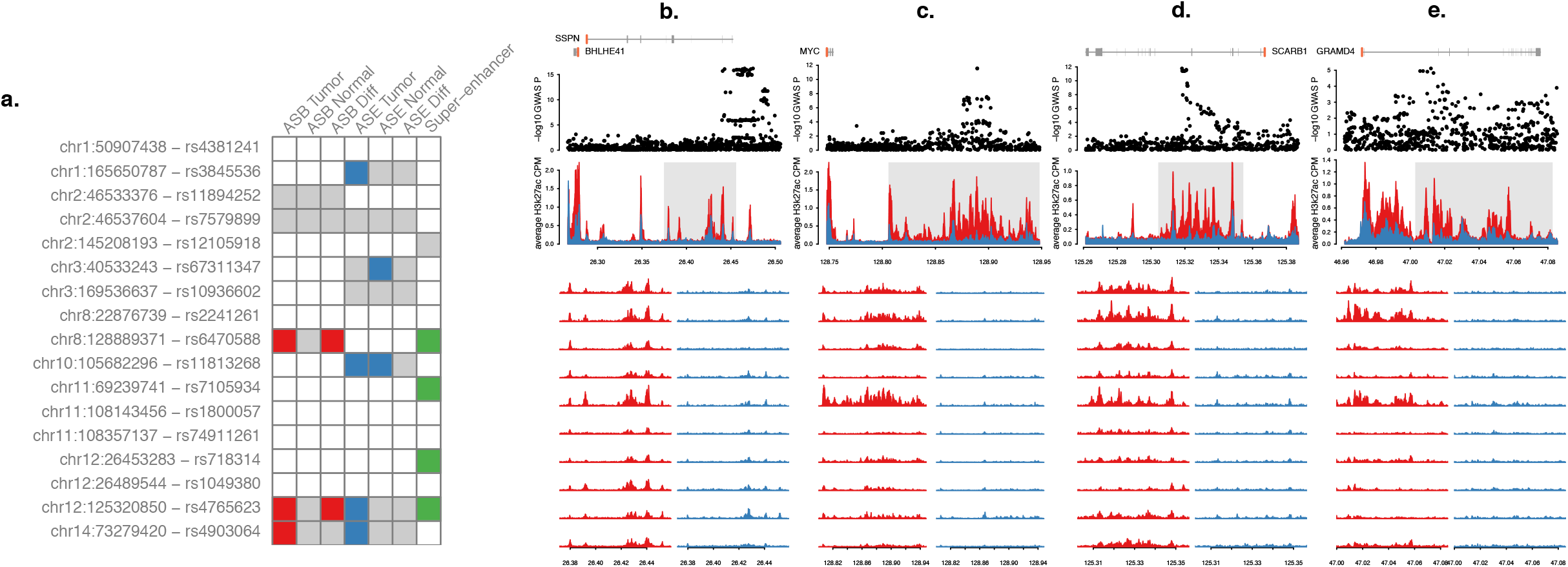
Allelically imbalanced functional features at RCC GWAS loci. (**a**) Each row lists the lead genome-wide significant SNP from *Scelo et al. 2017 Nat Comms* and each column corresponds to the presence of a functional feature. For ASB/ASE gray indicates sufficient heterozygous carriers to test for imbalance; for super-enhancer gray indicates the presence of an overlapping super-enhancer. Red indicates significant allele-specific binding (ASB) in H3k27ac (at 5% FDR genomewide); blue indicates significant allele-specific expression (ASE) in RNA-seq; green indicates significant tumor/normal differential enhancer activity. (**b**), (**c**), (**d**), (**e**) Four highlighted GWAS loci with significant differential super-enhancer activity. Panels (top to bottom) are: (1) gene/exon positions; (2) GWAS Manhattan plot of the locus; (3) average H3k27ac activity in tumor (red) and normal (blue); (4-13) H3k27ac activity in each individual tumor (red, left) and normal (blue, right).

We highlight the 12q24 GWAS risk locus as exhibiting all of the above functional qualities. The locus contains a super-enhancer that is tumor specific across all 10 individuals (**Figure 3d**). The lead GWAS risk variant is a significant T-asQTL (Q_tumor_=9.7×10^−7^) and d-asQTL (Q_d_=6.3×10^−3^) for this super-enhancer, as well as a significant T-asQTL for expression of *SCARB1* (Q=1.1×10^−53^) (**Figure 4**). The same SNP was a significant tumor eQTL (P=2.7×10^−15^ in all tumors; P=4.0×10^−4^ in subset that had matched normals) but had no effect in normal (P=0.62). Analysis of the tumor H3k27ac data revealed a high correlation between the genotype and overall activity (R2=0.78, p=0.007). Recent work has short-listed *SCARB1* as a transcriptional target of the hypoxia-inducible factor (HIF) which is suspected to play an important role in lipid metabolism and RCC ^28–31^. *SCARB1* encodes the SR-BI receptor which has been shown to be a of importance in ccRCC patients in 1 study ^32^ and previously for other cancers ^33^. In that ccRCC study, high expression of SRBI was associated with an unfavorable outcome. Furthermore, experimental knockdown of SRBI in RCC cell lines was shown to attenuate tumor proliferation and have downstream effects on HDL uptake and metabolic pathways ^32, 34^.

**Figure 4.**
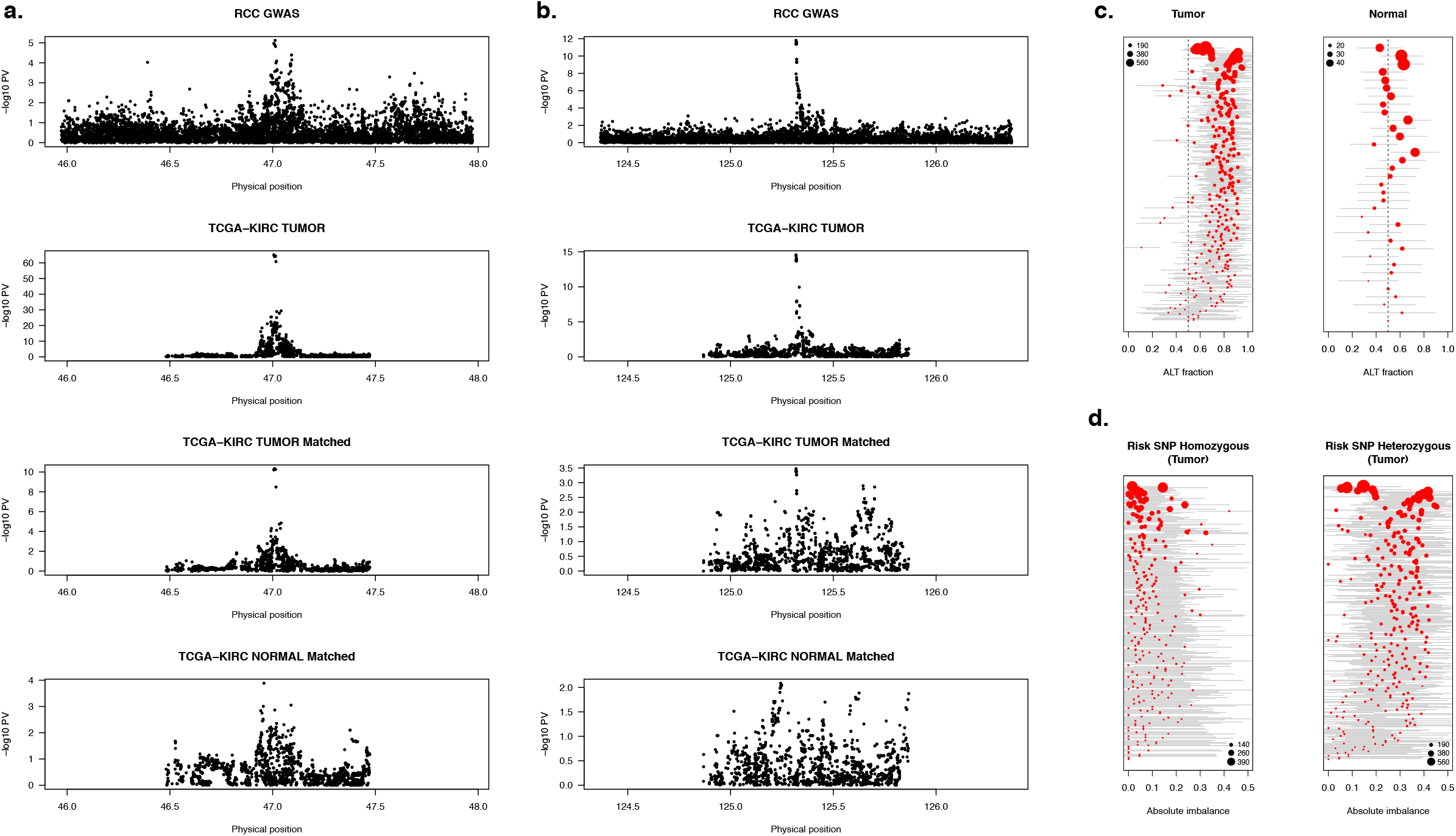
QTL signal at selected RCC GWAS loci. (**a**) Manhattan plots for SCARB1 locus (top to bottom): (1) GWAS association significance; (2) eQTL in all tumors; (3) eQTL in subset of 66 tumors with matched normals; (4) eQTL in 66 matched normals. Risk SNP is a nominally significant eQTL in the subset of tumors but not in the matched normals. (**b**) Manhattan plots for GRAMD4 locus as described in (**a**). Risk SNP is a highly significant eQTL in the subset of tumors but not in the matched normals. (**c**) Per-individual distribution of allelic imbalance for *GRAMD4* in heterozygous GWAS risk variant carriers. Each point represents an individual allelic-fraction (x-axis, log scale) in tumor (left) and normal (right). Point size corresponds to number of reads at that heterozygous haplotype. Gray line shows approximate standard error estimated likelihood ratio test. (**d**) Per-individual absolute tumor allelic imbalance at exonic SNP stratified by homozygous risk carriers (left) and heterozygous risk carriers (right). Homozygous risk SNP carriers are typically not significantly imbalanced, consistent with no secondary imbalance signal. Heterozygous risk SNP carriers are typically imbalanced, consistent with being the causal regulatory variant.

We further reasoned that overlap with an asQTL can increase the prior association for GWAS variants just below genome-wide significance and looked for colocalization with sites that were nominally significant (P<5×10^−7^) in the RCC GWAS. Indeed, 8/15 such variants showed overlap with asQTL peaks or genes (**Supplementary Table S5**). Notably, one SNP (rs714024) was significantly differentially imbalanced for chromatin (Q_tumor_=1.4×10^−7^; Q_d_=0.023), expression of the *GRAMD4* gene (Q_tumor_<1×10^−100^; Q_d_=1.9×10^−21^), and resided in a differentially active super-enhancer (**Figure 3e**). This SNP was a highly significant *GRAMD4* eQTL in tumor (P=3.5×10^−65^) but not in normal (P=0.01) and consistently imbalanced across the tumor (but not normal) heterozygous carriers (**Figure 4b,c,d**). Although rs714024 was not reported as a hit in the RCC GWAS, it was genome-wide significant (P=2×10^−8^) in the subset of clear-cell RCC cases, confirming its role as a germline risk variant ^1^. *GRAMD4* has been shown to modulate p-37 pathway induced apoptosis ^35^, making it a compelling example of a germline risk mechanism that interacts with the somatic environment (and thus primarily detectable in tumors). This is an instance where functional evidence prioritized a sub-threshold locus that would have otherwise been missed in the primary study.

We confirmed SE regulation for both *SCARB1* and *GRAMD4* by treating RCC cells with JQ1, a BRD4 inhibitor that preferentially affects SE-associated genes. Following 6 hours JQ1 treatment at the effective IC_50_ dose, *GRAMD4* showed a 33.6% (p-val = 8.48×10^−5^), 44.5% (p-val = 1.11×10^−4^), and 77.5% (p-val = 1.10×10^−4^) decrease in gene expression in 786-O, A498, and UMRC2 renal cell lines, respectively (**Supplementary Figure S8**). In comparison, *SCARB1* showed a decrease in expression of 8.91% in 786-O (p-val = 0.144), 62.6% in A498 (p-val = 2.83×10^−4^), and 28.8% in UMRC2 (p-val= 0.0177) (**Supplementary Figure S8**). We noted that both *GRAMD4* and *SCARB1* were generally more responsive to JQ1 than *MYC*, a known SE-regulated gene in many tumor types^23^. *GRAMD4* was particularly sensitive to JQ1 treatment across all cell types, and these data suggest that both genes are under control of cell-type specific super enhancers in ccRCC.

### Functional validation of *SCARB1*

We selected the *SCARB1* locus for further functional validation. Joint fine-mapping of the eQTL and GWAS signal identified 39 variants in the joint 95% credible set, of which 27 were also inside the asQTL peak (**Supplementary Table S6**). To further refine this variant set, we used accessible chromatin and HIF binding sites. These criteria identified a peak that contained two variants, rs4765623 and rs4765621, which are in perfect LD (**Figure 5a and 5b**). Experiments were performed in the 786-O cell line because it is heterozygous for these variants (**Supplementary Figure S9**). We selected rs4765623 as a reporter SNP for the regulatory region; 786-O is triploid and the rs4765623 genotype is CTT (C is the risk allele) (**Supplementary Figure S9a**). We also identified a heterozygous reporter SNP (rs5888) located in exon 8 of SCARB1 enabling measurement of AI in the *SCARB1* transcript (**Supplementary Figure S9b**). As expected, the allelic ratio was skewed and showed significantly higher contribution of the risk allele at both variants at the cDNA level (**Supplementary Figure S9c and 9d**). Next, allele specific enhancer activity was confirmed performing H3K27ac chromatin immunoprecipitation (ChIP) followed by Sanger sequencing. While the input conferred higher representation of the T allele, the IP sample was enriched for the C allele consistent with higher enhancer activity at the risk allele (**Supplementary Figure S9e**).

**Figure 5.**
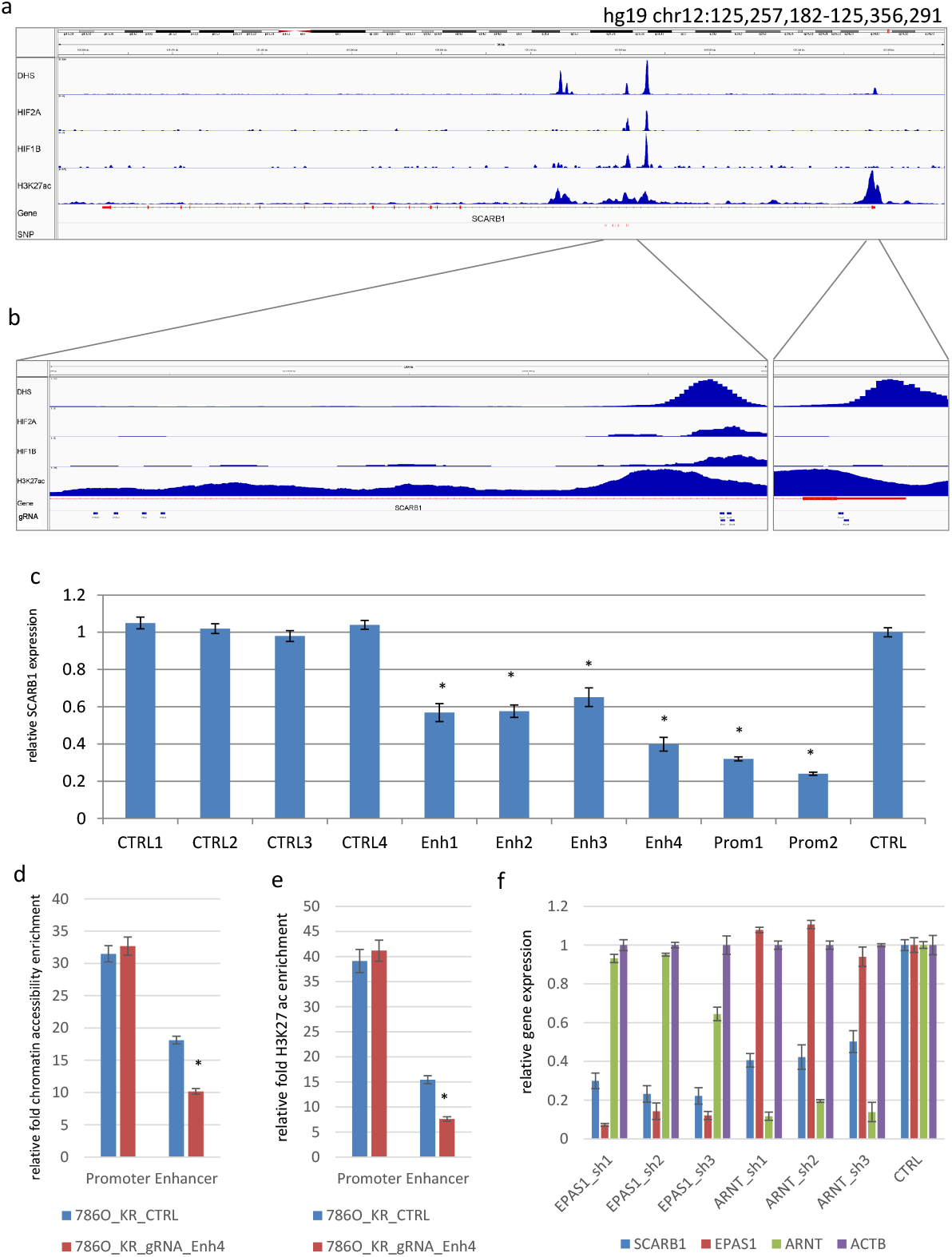
Enhancer activity and cell line genotypes at SCARB1 enhancer. (**a**), (**b**) and (**c**). To assess the impact of the candidate causal variants on SCARB1 expression, we performed CRISPR interference (CRISPRi) and designed four sgRNAs (Enh1-Enh4) against the rs4765623 region. Compared to the control gRNA, all four sgRNA significantly suppressed SCARB1 expression. Two sgRNAs (Prom1 and Prom2) were used as a positive control to directly target the SCARB1 promoter. As expected, these sgRNAs also suppressed SCARB1 transcript levels. To ensure that suppressing the intron did not affect SCARB1, we designed four sgRNAs (Top1-Top4) against candidate causal kidney SNPs that were contained within the same intron, but were located approximately 2.5 kb away from Enh1-Enh4 and did not overlap open chromatin. CRISPRi against this region did not alter SCARB1 expression. (**d**) Chromatin accessibility signals at SCARB1 enhancer position and promoter region were determined by ATAC-qPCR in 786-O cells. We used the top-scoring Enh4 sgRNA for this experiment. Results are expressed as relative enrichment normalized to non-accessible control genomic region. (**e**) H3K27Ac signals were determined by ChIP-qPCR in 786O cells. Results are expressed as relative enrichment normalized to non-acetilated control genomic region. (**f**) In order to identify the most likely trans- acting factor binding to the *cis*-element, potential regulatory TFs were targeted by shRNAs. Scrambled shRNA as a control was applied and relative gene expression was measured by qPCR. Suppression of ARNT and EPAS1 mRNA levels with three different shRNAs demonstrated significantly reduced SCARB1 expression indicating that these TFs regulates the SCARB1 gene via this enhancer. Data represent the average and standard deviation of 3 biological replicates (C-F). Each measurement was performed in triplicates. Significance determined by Student’s t test. *p < 0.05.

In order to evaluate enhancer activity and its regulatory effect on *SCARB1* expression, we performed CRISPR interference (CRISPRi) using the dCas9-KRAB repressor construct ^36^. Four single guide RNAs (sgRNAs) targeting the DNase peak demonstrated strong suppression of *SCARB1* transcript levels (**Figure 5c**). As a stringent negative control to determine *SCARB1* specific enhancer activity, we targeted the top two (rs10846748 and rs10846750) most significant SNPs from the joint fine-mapping by 4 sgRNA (Ctrl1-4), which reside in the same intron and super enhancer but do not exhibit TF/DHS activity. Targeting these SNPs had no effect on *SCARB1* gene expression as expected (**Figure 5c**). These data demonstrate that this peak is a functional *SCARB1* enhancer. To evaluate the impact of CRISPRi on chromatin accessibility and H3K27ac signal, we performed ATAC- and H3K27ac ChIP-qPCR. Both ATAC-seq and H3K27ac signals demonstrated significant decrease at the enhancer position (**Figure 5d and 5e**).

Prior ChIP-seq data showed binding of multiple TF binding sites at this locus (**Figure 5a**), leading us to performed RNAi experiments to suppress the upstream TFs. ShRNAs were designed against *ARNT* (*HIF1B*) and *EPAS1* (*HIF2A*) and transduced into 786-O. Suppression of these factors demonstrated decreased *SCARB1* expression implying that HIF1B and HIF2A regulate *SCARB1* through this enhancer (**Figure 5f**).

### d-asQTLs are enriched for GWAS risk heritability

To investigate the relationship between asQTLs and RCC risk more broadly, we integrated our analysis with the most recent RCC GWAS of 10,784 cases and 20,406 controls 1 and applied stratified LD-score regression (S-LDSC ^10^) to estimate risk heritability residing in relevant functional features. We observed enormous heritability enrichments for H3K27ac asQTL peaks, larger than all classes of non-asQTL peaks and asQTL genes: d-asQTLs covered 0.7% of the genome while explaining 23% of the overall RCC heritability (32x enrichment; P=0.002); T-asQTLs covered 1.4% of the genome while explaining 48% of RCC heritability (28x enrichment; P=6.6×10^−05^); with a similar, although non-significant, enrichment for N-asQTLs (**Figure 6a, Supplementary Table S7**). These asQTL enrichments were ~2.8x higher than enrichments for the full set of H3K27ac peaks detected in tumor (16x; P=1×10^−9^) or normal kidney tissue (14x; P=8×10^−7^) showcasing H3K27ac activity as a key functional feature for RCC risk and allelic imbalance as a means to further refine these features. In contrast, super-enhancers, which have been posited as important drivers of cancer, were significantly less enriched (5x; P=6×10^−6^) suggesting lower relevance for germline risk mechanisms. Indeed, super-enhancers that were active/gained in tumors were the only functional category to have no marginally significant enrichment. To evaluate whether the d-asQTL enrichment was specific to RCC (rather than, for example, tagging regulatory regions of general importance to complex traits) we performed the same analysis across a collection of 95 publicly available GWAS traits 1037. The average enrichment for d-asQTL peaks across these generic traits was 3.4x, with no single trait approaching the 32x enrichment observed for RCC (see **Supplementary Table S8, S9, S10** for trait-specific results).

**Figure 6.**
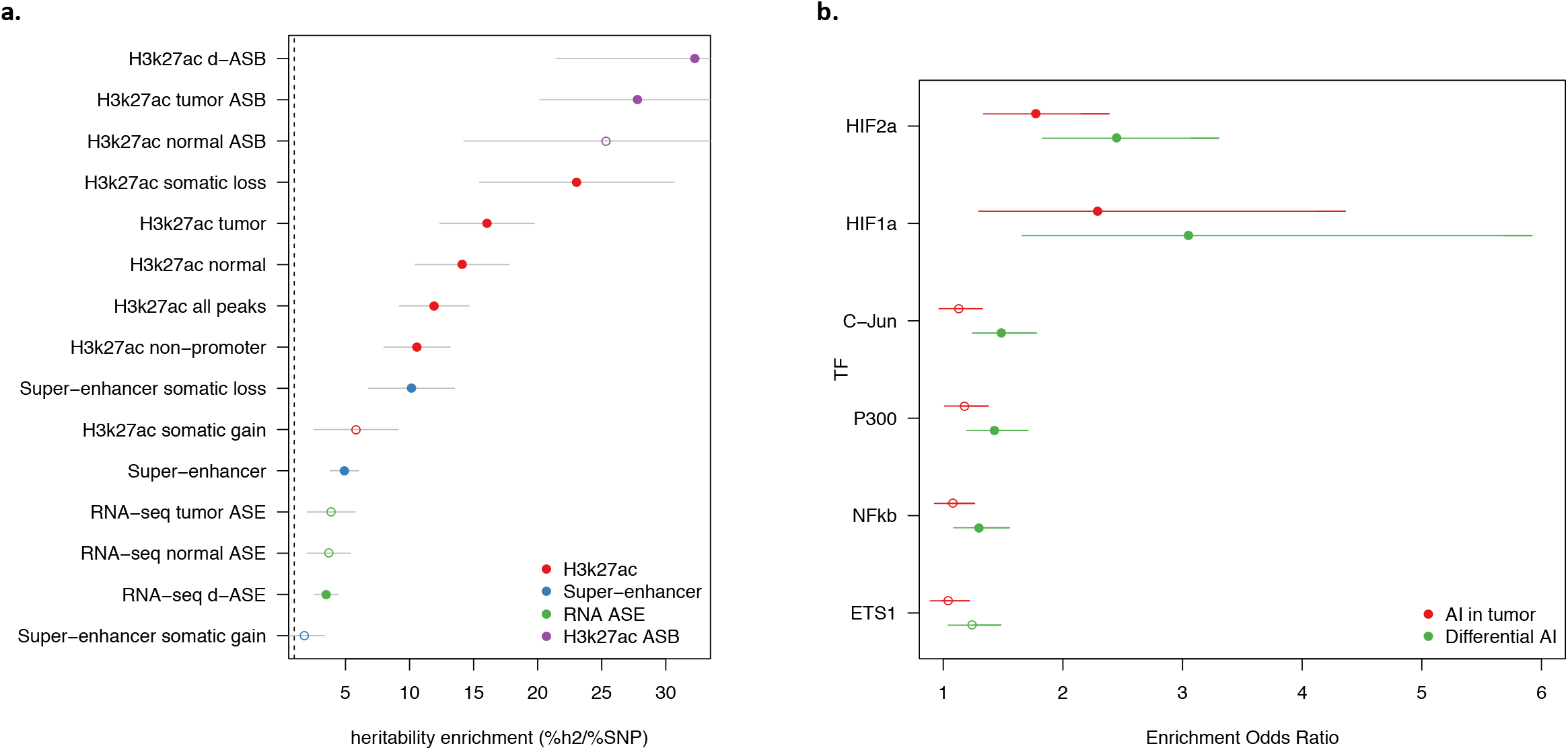
Heritability and functional enrichment across imbalanced features. (**a**) For each functional feature (y-axis) the heritability enrichment (x-axis; fraction of heritability / fraction of SNPs) is shown with points and standard error bar. Filled points indicate statistical significance after correcting for number of annotations tested. Colors indicate feature class shown in legend. Dashed line at 1.0 (no enrichment). (**b**) Enrichment of TF binding in tumor imbalanced peaks relative to normal imbalanced peaks. Each point report the odds ratio of a tumor asQTL peak (red) or d-asQTL peak (green) overlapping a given TF relative to normal asQTL peaks (Fisher’s exact test). Error bars report the 95% confidence interval around enrichment. Solid points indicate significant enrichment after Bonferroni correction for 6 tests.

We next tested an annotation corresponding to genes with significant RNA-seq asQTLs. We defined 100kb windows around the TSS of asQTL genes to match the discovery window and then chose the 200 most significant asQTL genes for each state, yielding an annotation covering ~1.8% of the genome (comparable to ~1% of the genome covered by the imbalanced peak annotations). As with the peak enrichment, we selected regions rather than individual QTLs due to the recent finding that lead eQTLs exhibit atypical LD characteristics that can bias enrichment estimates ^38^. We observed significant enrichment only for the d-asQTL genes (3.5x; P=8×10^−4^), which was still substantially weaker than the enrichment at imbalanced H3K27ac peaks (**Figure 6a, Supplementary Table S7**). This difference in enrichment is stark and suggests that H3K27ac asQTLs are more precisely localizing causal germline risk variants than overall expression asQTLs. We further intersected the asQTL gene annotations with H3k27ac peaks, which increased the resulting enrichments but did not exceed the enrichment from imbalanced peaks alone (**Supplementary Figure S10, Supplementary Table S7**). The lower enrichment at asQTL genes is therefore not due to the broad region definition, but suggests that the strongest asQTL genes are prioritizing generic cell processes that are less relevant to disease risk than asQTL peaks. In sum, these findings reveal d-asQTL H3k27ac peaks as features that harbor the greatest per-variant effect in RCC GWAS and represent important novel targets for disease fine-mapping and functional validation.

### d-asQTLs are enriched for tumor-specific TF activity in the hypoxia pathway

We hypothesized that the H3k27ac T-/d- asQTLs may be explained by emergent binding of tumor-centric TFs at regulatory variants that are unbound and balanced in the normal state. Under this model, we would expect to see an enrichment of TF activity at d-asQTL peaks relative to active peaks without asQTLs (which reflect trans effects) or with N-asQTLs (which reflect general regulatory processes in renal cells). We focused on six TFs recently investigated by 39 in RCC cell lines and shown to be enriched near tumor-specific enhancers. First, we used the TF ChIP-seq from 39 to confirm that all six TFs had more binding sites in tumor-specific enhancers relative to all enhancers in our samples, with the strongest enrichment observed for HIF1A binding in distal tumor-specific enhancers (Fisher’s exact OR=10.6, P=3×10^−49^; **Supplementary Table S11**). Next, we sought to quantify enrichment of each TF at asQTLs while controlling for the fact that QTL detection power is higher in more active peaks (which contain more allelic reads). For each set of T-/N-/d- asQTL peaks and each TF we tested the number of overlapping TF binding sites against a “background” set of random H3K27ac peaks that were matched to have the same coverage distribution at heterozygous sites (see Methods). All six TFs exhibited more binding sites at asQTL peaks in all three states, but HIF1A/HIF2A specifically exhibited greater enriched in T-/d- asQTLs than N-asQTLs (Fisher’s exact P<2×10^−12^; **Supplementary Table S12**). Directly comparing T-/d- asQTLs to exclusively N-asQTLs confirmed significantly more HIF1A/HIF2A binding sites in the former (**Figure 6b, Supplementary Table S13**). In both comparisons the enrichment for d-asQTLs was greater than for T-asQTLs. These enrichments support a model of widespread germline variants interacting with tumor TF binding as we had observed functionally for the *SCARB1* locus.

Motivated by the prioritization of HIF1a/HIF2a binding sites, we searched for SNPs in the RCC GWAS that overlapped asQTL peaks and HIF1a/HIF2a active sites (or sites that disrupted known HIF motifs, see Methods) and identified a total of 8 SNPs where the GWAS association was significant after Bonferroni correction: 3/8 loci were identified or connected to target genes or enhancers from asQTLs in this study: *DPF3, SCARB1*, and *GRAMD4;* another 4/8 loci were previously linked to HIF binding and/or experimentally supported: *EPAS1* (HIF2a), *CCND1, BHLHE41*^40^, and *MYC* ^41^ (**Figure 3b,c, Supplementary Table S14**) and the remaining SNP was in an intron of the *RBPMS* gene but could not be connected to a target. In total, asQTLs were observed at all four loci previously implicated through HIF binding and highlighted three novel gene targets at four loci without previous experimental follow-up.

## DISCUSSION

As GWASs have implicated thousands of disease-associated loci, many approaches have emerged for linking these associations to their target genes. The availability of population-scale epigenomic data further empowers discovery of cis-regulatory elements that may mediate between the risk variant and target gene and thus constrain putative targets for experimental validation.

Here, we propose a novel method that leverages allelic imbalance to identify *cis*-regulatory elements while accounting for SCNA in cancer samples. The robustness of the approach additionally allows us to test for changes in *cis* regulation between tumor and normal tissues. Application of our method to ChIP-seq and RNA-seq data from tumor and normal samples provided multiple insights into polygenic regulatory mechanisms. First, germline allelic imbalance in binding and expression is ubiquitous in the tumor and often significantly different from the matched normal. These differences were significantly but only partially explained by local SCNA loss events and were not significantly associated with SCNA gains. Second, enhancer peaks exhibiting tumor-specific imbalance were enriched for binding of tumor-specific TFs in the HIF pathway. HIF pathway disruptions have been previously implicated in both the germline and somatic etiology of RCC. We thus postulate a model where certain germline variants are altering inactive TF binding sites in the normal cell (with corresponding allelic balance of the regulatory element) and then become active regulatory variants in response to tumor-centric TF activity after transformation. Third, we observed that differentially imbalanced enhancer peaks are slightly more enriched for cancer risk heritability than imbalanced enhancers in the normal tissue, and substantially more enriched than peaks that did not exhibit allelic imbalance. This is consistent with our model that differentially imbalanced peaks prioritize cancer relevant TF binding sites. Finally, we identified three GWAS loci for which the lead risk SNP was a tumor-specific d-asQTL for an overlapping chromatin peak, of which two were also tumor-specific d-asQTLs for a nearby gene. We experimentally validated the *SCARB1* locus through CRISPRi of the implicated regulatory element and also confirmed an association with upstream HIF1B/HIF2A activity, consistent with our germline-somatic interaction hypothesis.

Our study has several limitations. First, the chromatin data evaluated here is likely capturing only a small fraction of all cis-regulatory elements, which can be expanded through increased sample size and a broader set of epigenomic marks. In particular, fewer than half of the active chromatin peaks carried sufficient heterozygous variants and allelic reads to be tested in our data. In contrast, the large sample size available for tumor RNA-seq allowed us to identify significant asQTLs for >60% of tested genes. Second, we considered chromatin asQTLs that resided inside enhancer peaks for simplicity of interpretation, but larger sample sizes will allow fine-mapping of causal regulatory variants that lie outside the peak or that have complex multi-peak influences ^42^. While allelic imbalance was highly effective for discovery of regulator elements, we found that fine-mapping of the causal variant still required traditional QTL analyses and larger sample sizes due to the difficulty in modelling allelic imbalance haplotype structure. Third, we did not have the power to distinguish imbalanced features driven by local SCNA events (which we estimate to explain 14-16% of the T-/d- asQTLs). We caution that asQTLs observed in a small number of individuals could thus be driven by SCNAs rather than germline polymorphisms and recommend leveraging large-scale QTL studies for confirmation. Lastly, as with all such integrative analyses, the co-occurrence of imbalanced chromatin/expression features and GWAS risk variants does not guarantee a direct causal relationship. Other plausible mechanisms include: incidental tagging of multiple independent causal variants; independent pleiotropic effects on the two features by the same causal variant; or an indirect causal relationship mediated by some other gene or regulatory feature that was not observed in this data.

A model where tumor-specific activity is as relevant to cancer risk as normal activity is intriguing and highlights an often-overlooked source of mechanistic information for GWAS. Regulatory sites that increase risk may do so through mechanisms that are positively selected in the evolving cancer population, and thus be more detectable in tumor tissues. Alternatively, the normal tissue may be more cell-type heterogeneous and thus obscure the precursor cells; whereas the tumor is typically more clonal and thus more reflective of precursor activity. d-asQTLs in tumor may thus be highlighting precursor vs. non-precursor regulatory differences rather than strictly tumor-specific activity. Precise measurements of cell-composition or single-cell transcriptomics may soon empower us to evaluate these hypotheses directly. The co-occurrence of tumor-specific activity and risk heritability is not limited to our study but has been previously observed in RCC for HIF binding ^41^; prostate cancer for androgen receptor binding ^20^; and breast cancer for estrogen and FGFR2 signaling ^43^. Together, these findings suggest general tumor-centric mechanisms may be relevant for cancer risk and motivate further investigation of molecular measurements from tumor and normal samples.

## METHODS

### Sample preparation

Specimens were obtained from patients, with appropriate consent from the institutional review board and were previously submitted to TCGA ^44^. Tissue ChIP was performed as previously described ^45^. Briefly, one frozen 2 mm core was pulverized in the pulverization bag using the Covaris CryoPrep system (Covaris, Woburn, MA) once at intensity 3 and once at intensity 4. The tissue was then fixed using 1% formaldehyde (Thermo fisher, Waltham, MA) in PBS for 10 minutes at RT with rotation, and quenched with 125 mM glycine for 10 min at room temperature with rotation. After rinsing with ice-cold PBS twice, chromatin was resuspended and lysed in ice cold lysis buffer (50mM Tris, 10mM EDTA, 1% SDS with protease inhibitor) for 10 minutes. Chromatin was sheared to 300–500 base pairs using the Covaris E210 sonicator (AFA: 5% duty cycle, 5 intensity, 200 cycles/burst) for 10 min. 1% of chromatin was saved as input for each sample. 5 vol of dilution buffer (1% Triton X-100, 2 mM EDTA, 150 mM NaCl, 20 mM Tris HCl pH 8.1) was added to the rest of chromatin and then, the sample was incubated with 1ug H3K27Ac antibody (DiAGenode, C15410196, Denville, NJ) coupled with protein A and protein G beads (Life Technologies, Carlsbad, CA) at 4 degree overnight. The chromatin was washed with RIPA washing buffer (0.05M HEPES pH 7.6, 1 mM EDTA, 0.7% Na Deoxycholate, 1% NP-40, 0.5M LiCl) for 5 times each for 10 minute and, rinsed with TE buffer (pH 8.0) once. The sample was resuspended in elution buffer (50mM Tris, 10mM EDTA, 1% SDS), treated with RNase for 30 minutes at 37 degree, and incubated with proteinase K for overnight at 65 degree. Sample DNA as well as its input were extracted using Qiagen Qiaquick columns, and were prepared as the sequencing libraries using the ThruPLEX-FD Prep Kit (Rubicon Genomics, Ann Arbor, MI). Libraries were sequenced using 75-base pair single reads on the Illumina platform (Illumina, San Diego, CA) at the Dana-Farber Cancer Institute.

### H3k27ac peak calling

H3k27ac peaks were called from mapped reads using MACS2 (v2.1.1.20160309) with default settings in broad peak mode ^46, 47^. Super-enhancers were called from H3k27ac peaks, restricted to non-promoters, using the ROSE pipeline (v0.1) ^23, 48^. DNA input for each sample was used as a control for all calls.

The edgeR software was used to test for differences in tumor/normal peak activity using the exact negative-binomial model for mean group differences ^49^. Overdispersion was estimated for each peak separately (i.e. “tagwise”) using empirical Bayes.

Unsupervised clustering was performed by quantifying the ChIP-seq read activity in each peak/sample, estimating a pairwise correlation matrix across all samples, and performing agglomerative hierarchical clustering using the complete linkage method (implemented in the R hclust function).

Tumor/normal expression and methylation quantifications were downloaded from TCGA Firebrowse and assigned to the nearest chromatin feature.

### Genotype quality control and phasing

SNP genotype calls using Birdsuite ^50^ were downloaded from the TCGA legacy archive and phased/imputed using the EAGLE pipeline provided by the Michigan imputation server ^51^. Eagle reports typical phasing accuracy of >1MB so no additional modeling of phase uncertainty was performed. Imputed SNPs were retained if they had INFO>0.9; locus missingness <5%; hardy-weinberg equilibrium p-value > 5×10^−6^ (thresholds based on GTEx recommendations ^14^). Variants of any frequency were retained for the ChIP-seq analysis; and variants with MAF>1% in the KIRC cohort were retained for the RNA-seq analysis. Birdsuite tumor/normal CNA calls were downloaded from TCGA Firebrowse.

### ChIP-/RNA-sequence read quality control

Read mapping bias is a particular concern for measuring AI and we ran all RNA-seq and ChIP-seq data through the WASP re-mapping pipeline ^17^. Briefly, for every read within a heterozygous site, a synthetic read is generated carrying the alternative allele and re-run through the full mapping pipeline. Only reads with synthetic partners mapping unambiguously to the same site are retained for subsequent analysis. After re-mapping, duplicate reads are dropped randomly to avoid allele-specific biases. Finally, read counts for each site and allele were computed using GATK and the ASReadCounter module ^26^. Our computational pipeline is publicly available (see Data Availability).

### Stratified Allele-Specific (stratAS) test

We propose a statistical framework to identify allelic imbalance and tumor/normal differential imbalance. A key novelty of our approach is that it directly models tumor-specific confounders and evaluates differential imbalance under a formal likelihood model. We derive the imbalance statistic for a single individual and site; then extend it to analysis of multiple sites within the population; and close with a comparison to traditional QTL-based interaction tests.

#### Allele-specific test at a single site

Rather than using between-sample differences in *y*, we propose to test heterozygous germline variants for significant imbalance of reads favoring one allele *within* each individual – thus circumventing environmental confounding. The key insight of this approach is that any trans-acting confounders will impact reads for each allele independently and equally (in expectation), and thus not bias the mean estimate of imbalance. For a given heterozygous germline SNP *j* in individual *i* we model the tumor/normal reads overlapping the site (*T, N* respectively) as coming from a beta-binomial distribution: *T_alt, i_*|*T_ref,i_*^~^ *BetaBin*(π_*T,j*_, ρ_*T, ij*_); *N_alt,i_*|*N_ref, i_*^~^ *BetaBin*(π_*N, j*_, ρ_*N, ij*_), where [π_*T*_, π_*N*_] are the mean allelic ratios and [ρ_*T, ij*_, ρ_*N, ij*_] are locally defined, per-individual sequence read correlation parameters.

We test for T/N differential imbalance by likelihood ratio test between the models π_*N,j*_ = π_*T, j*_ and π_*N, j*_ ≠ π_*T, j*_, maximizing the likelihood of each model by a standard 1-dimensional optimization.

To account for over-dispersion due to somatic structural variations, the ρ_*, *ij*_ parameters are estimated for each individual from all heterozygous read-carrying SNPs within the overlapping region of consistent copy number. We observed empirically that reads from tumor sequencing are highly over-dispersed and correlated with local somatic copy number, which we account for using this CNA-specific parameter.

#### Allele-specific test incorporating haplotype information

We leverage high-quality haplotype phasing^51^ to further extend this test to multiple variant analysis and fine-mapping. First, reads from all heterozygous variants within a functional feature (chromatin peak or multiple exons, for example) can be aggregated together along the haplotype under the assumption that the regulatory effect is consistent across sites. As H3k27ac peaks tend to be broad this approach significantly increases power by aggregating read coverage across multiple markers. Second, any heterozygous variant outside the chromatin peak can still be tested for imbalance by assigning each allele carrier the reads on their corresponding haplotype. For a single individual this is mathematically identical to testing the peak SNPs, but within the population this allows one to resolve causal regulatory SNPs that are distal to the chromatin activity.

### Simulation framework

We sought to evaluate our method under realistic tumor/normal read distribution conditions, as this has the biggest impact on AI approaches. We based our simulations around the real peaks, genotypes, and coverage counts observed in the H3k27ac samples, as well as the corresponding CNA-specific over-dispersion parameters. For a given allele fraction, we sampled reference/alternative reads at each site and individual from a beta-binomial distribution based on these parameters. We then evaluated each method across all peaks and sites in the study to mimic the full coverage distribution. Under the null, we simulated a tumor and normal allele-fraction of 0.5. Under the alternative, we simulated a tumor allele-fraction ranging from 0.6-0.9 (in increments of 0.1) and tested for imbalance in the tumor, as well as tumor/normal difference in imbalance. We evaluated the binomial, global beta-binomial, and CNA-specific (“local”) beta-binomial model across these scenarios. For the differential simulations, we additionally evaluated the Pearson correlation and c-ASE Z-score approximations (see **Supplementary Note**).

### stratAS parameter setting and estimation

For analysis of real data, we used slightly different test inclusion criteria for the ChIP-seq and RNA-seq data due to their vast differences in sample size and read distributions.

For ChIP-seq peaks, we tested only variants within the peak as this sample size was insufficient to perform thorough fine-mapping. We restricted tests to peaks with at least 5 reads in total and at least one read supporting each haplotype.

For RNA-seq exons, we tested all common variants within 100kb of the TSS (having been shown to harbor the majority of regulatory variation in eQTL analyses). Sites were only tested if they had >1% heterozygous genotype carriers, and only heterozygotes with at least 5 reads supporting both alleles were evaluated.

A local beta-binomial overdispersion parameter ρ_*, *ij*_ was fit for each individual and region of contiguous copy number, using the reads at all contained heterozygous sites. Inference was performed using the vector generalized linear model in the VGAM R package.

### eQTL analyses

A traditional eQTL analysis was carried out for comparison of detection power between eQTL and asQTL approaches. Genotypes underwent the same QC process as the gene asQTL analysis and variants within 100kb of the TSS of each gene were again tested for association to overall gene expression. 10 principal components were computed from the gene expression matrix (for tumor and normal samples separately) and included as covariates to account for trans variation. Association was performed using fastQTL ^52^.

### Regression model quantifying contribution of local features to asQTLs

For each sample and state we restricted to peaks with >50 allelic reads and identified asQTLs using stratAS. Each tested peak was then labeled with a 1/0 for imbalance/balance in a sample, as well as the corresponding local features in that sample: (i) significant CNA loss, (ii) significant CNA gain, (iii) absolute CNA copy change from diploid, (iv) coverage, (v) N-asQTL in the matched normal. A logistic regression model was then used to jointly link the local features to imbalance across all peaks and individuals. The marginal variance explained by each feature was also quantified using Nagalkerke R2 with only that feature in the model.

### Estimating cross-state asQTL concordance using bootstrapped FDR

We quantified the fraction of asQTLs detected in one state that replicated in another state using the Storey pi0 statistic, which estimates the faction of null associations from a p-value distribution ^53^. Statistics were estimated using the bootstrapped method over increasingly stringent discovery FDR thresholds until the replication estimate plateaued.

### Partitioning GWAS heritability

We applied stratified LD-score regression (S-LDSC) to estimate feature-specific enrichment of association statistics ^10^. In brief, S-LDSC uses the association strength and LD to variants in a given functional category (such as a relevant set of enhancers) to infer the total *causal* contribution of those variants towards risk. This approach has been shown to be highly powerful even when no individual variants are significant, by aggregating enrichment across the full polygenic spectrum of variants.

### Enrichment for eQTLs from GTEx at asQTLs

We quantified the overlap of each of the T-/N-/d- asQTLs with eQTLs from the GTEx Consortium ^14^ across all tissues. For each set of asQTL peaks, we computed the fraction of SNPs in the peaks that were also GTEx eQTL in the given tissue. We performed the same computation for the same number of randomly sampled non-asQTL peaks (peaks that met all criteria for asQTL testing but did not have significant imbalance in that state), resampling 1000 times to produce a background distribution and compute confidence intervals.

### HIF binding and motif analysis

TF peak calls were downloaded from ChIP-seq analyses performed by ^39^. For each T/N/d state, enrichment odds ratios was evaluated by a Fisher’s exact test comparing the number of asQTL peaks overlapping TF peaks (i.e. the odds that an asQTL peak contains a TF) versus the number of TF peaks overlapping the “background” peaks in the same state (i.e. the odds that a background peak contains a TF). We then considered three definitions for background peaks: (i) all peaks that could be tested for asQTL (i.e. contained heterozygous variants with allelic reads) and were non-significant; (ii) peaks from (i) that were additionally subsampled to match on quintiles of the coverage distribution for the foreground asQTL peaks; (iii) a set of significant N-asQTL peaks that were exclusive to the N state. Separately, asQTL and GWAS variants were assessed for motif disruption of the HIF family using the *motifbreakr* software ^54^.

### Renal cell carcinoma (RCC) cell lines

The A-498 cell line was obtained from ATCC and maintained in DMEM (Invitrogen) supplemented with 10% Fetal Bovine Serum (FBS). The 786-0 RCC cell line was obtained from ATCC and maintained in RPMI-1640 (Invitrogen) supplemented with 10% FBS. The UM-RC-2 cell line was obtained from ECACC (Sigma) and maintained in DMEM supplemented with 1% Non-Essential Amino Acids with 10% FBS. All media was supplemented with 100 U/mL penicillin, and 100 mg/mL streptomycin (Invitrogen) and all cell lines were cultured at 37°C with 5% CO2. Each line was authenticated by short tandem repeat profiling and tested negative for mycoplasma (ABMG) in every month.

### Lentivirus production and establishment of stable cell lines

HEK293T cells (Sigma) were cultured in DMEM supplemented with 10% FBS. For lentivirus production transfer vectors and packaging mix (Cellecta, cat.no.: CPCP-K2A) were delivered by BTX (Harvard Apparatus) electroporation. To generate stable 786-O-dCas9-KRAB expressing cell line the Lenti-dCas9-KRAB-blast (Addgene #89567) was used. Virus transduced cells were maintained for 2 weeks under blasticidin (10 μg/ml, Invivogen) selection. For stable sgRNA and shRNA expression 3 days puromycin (2 μg/ml, Invivogen) selection was applied.

### Measure of allelic imbalance

PCR products were generated from gDNA, chromatin immunoprecipitation DNA and heteronuclear cDNA and subjected for Sanger sequencing (Genewiz).

### Design and creation of gRNA constructions

SgRNAs were designed using a previously published algorithm ^55^. For CRISPRi experiments, gRNA sequences were synthesized as complementary single stranded oligonucleotides with sticky ends (detailed protocol also available at https://media.addgene.org/data/plasmids/43/43860/43860-attachment_T35tt6ebKxov.pdf). gRNA target sequences are listed in **Supplementary Table S15**.

Following annealing, oligonucleotides were cloned into pXPR_BRD003 lentiviral gRNA expression vector by Golden Gate cloning. Stbl3 chemically competent cells (Invitrogen) were transformed with the gRNA constructs. Clones were verified by Sanger sequencing (Genewiz). Clones were propagated in LB broth under carbenicillin (100 μg/ml, Gibco) selection over night at 37°C and plasmid DNA was purified using QIAGEN Plasmid Plus Midi kit.

### Design and creation of shRNA plasmids

shRNAs were designed using the GPP web portal (https://portals.broadinstitute.org/gpp/public/) ^56^. Hairpin oligo nucleotides (Supplementary Table S15) were ordered from Integrated DNA Technologies and after phosphorylation and annealing were cloned into pLKO.1 - TRC shRNA cloning vector (Addgene #10878) into AgeI/EcoRI site. Clones verified by Sanger sequencing (Genewiz). Clones were propagated in LB broth under carbenicillin (100 μg/ml, Gibco) selection over night at 37°C and plasmid DNA was purified using QIAGEN Plasmid Plus Midi kit.

### Quantitative RT-PCR

Total RNA was isolated using NucleoSpin^®^ (Macherey-Nagel) RNA isolation kit. cDNA was synthesized from 1 μg total RNA using the High Capacity cDNA Reverse Transcription Kit (Applied Biosystems). Quantitative real time PCR was performed on Light Cycler 480 II instrument (Roche) using Power SYBR^™^ Green PCR Master Mix (Applied Biosystems). Oligo nucleotides were synthetized (Integrated DNA Technologies) with standard purification. Primer sequences used for qRT-PCR are listed in **Supplementary Table S15**. Relative gene expression was calculated based on the ddCt method. Each sample was measured by three biological and technical replicates, average and standard deviation were calculated. Statistical tests used and p-values are described in the figure legends.

### Histone ChIP-qPCR

H3K27ac ChIP-qPCR was performed as previously described ^57^. Briefly, 5 x 10^6^ 786-O cells were fixed using 1% paraformaldehyde for 10 minutes at room temperature and quenched with glycine for 5 minutes. Cells were resuspended in 1 ml shearing buffer (10 mM Tris pH 8.0, 5 mM EDTA, 0.1% SDS) and sonicated (Covaris) with the following settings (Peak Incident Power = 100 W, Duty Factor 5%, Cycles per Burst = 200 for 10 minutes) to obtain 200-800 bp range of average shared chromatin size distribution and centrifuged at 13,000 r.p.m. for 10 min at 4 °C. Immunopreciptation was performed overnight with 1 μg of antibody. Antibody used: anti-H3K27ac (Diagenode C15410196). Protein A/G magnetic beads (Life Technologies) were added and incubated for 2 hours at 4°C. Immunoprecipitates were washed with low salt buffer (20 mM Tris pH 8.0, 150 mM NaCl, 2 mM EDTA, 1% Triton), high salt buffer (20 mM Tris pH 8.0, 500 mM NaCl, 2 mM EDTA, 1% Triton), LiCl buffer (100 mM Tris pH 8.0, 500 mM LiCl, 1% NP-40, 1% deoxycholate), and twice with TE (10 mM Tris pH 8.0, 1 mM EDTA). DNA was eluted with SDS/NaHCO3 elution buffer (1% SDS, 0.1M NaHCO3) at 65°C. Crosslinking was reversed by treating with proteinase K at 65°C for 6 hours followed by RNase treatment for 30 min at 37°C. DNA was purified using QIAGEN PCR purification kit, and quantified using qPCR. Primers used are listed in **Supplementary Table S15**.

### Interrogation of chromatin accessibility (ATAC-qPCR)

Open chromatin regions were revealed by applying Omni-ATAC method ^58^, using 50 000 cells/samples. Briefly, cells were lysed in ATAC-Resuspension Buffer, after washing out lysis reagents, nuclei were collected and resuspended in transposition mixture containing Tn5 transposase enzyme. Following 30 minutes incubation at 37°C, DNA was extracted by using the Zymo DNA Clean and Concentrator-5 Kit (cat. no.: D4014), and used for amplification. Primers used are listed in **Supplementary Table S15**.

### JQ1 treatment of SCARB1 and GRAMD4

786_O, A498, and UMRC2 were seeded in 96-well plates at a density of 10,000 cells/well. The effective IC_50_ concentration was determined for each cell line following a 72-hour treatment with JQ1 (Tocris) at concentrations from 1.52 nM to 10 μM. Cell populations were measured using Cell Titre Glo (Promega), and dose response curves were generated using GRMetrics ^59^. The effective IC_50_ of the cell lines 786_O, A498, and UMRC2 was determined to be 461 nM, 752 nM, and 2177 nM, respectively. Cells were then treated with JQ1 at the effective IC_50_ value for 6 hours and RNA was harvested using NucleoSpin RNA Plus columns (Macherey-Nagel). RNA expression levels were quantified after reverse transcription with MMLV H-(Promega) using the TaqMan gene expression system (Thermo Fisher, probe numbers Hs00411719 and Hs00969821). Expression levels of genes were compared using the ddCT method, using Actin B (Hs01060665) and GAPDH (Hs02786624) as the control genes. A non-targeting DMSO vector negative control was used to compare the relative gene expression as a result of JQ1 treatment no inhibition. DMSO and JQ1 treatment were performed in technical duplicates for three biological passages.

### Data/Software availability

The stratAS software and asQTL pipeline is available at https://github.com/gusevlab/stratAS. H3k27ac reads, peak calls, and allele-specific calls will be deposited into appropriate databases prior to publication.

## ACKNOWLEDGEMENTS

We would like to thank Stephen Chanock, Simon Gayther, Bryce van de Geijn, Alexander Gimelbrant, and Bogdan Pasaniuc for helpful discussions. We would also like to thank Ghislaine Scelo, Mark Purdue, Stephen Chanock, and the renal cell carcinoma GWAS consortia for the RCC GWAS association statistics.

AG was supported by R01-CA227237 and the Claudia Adams Barr Foundation Award. K.L. is supported in part by a Liz Tilberis Award from the Ovarian Cancer Research Alliance (OCRA, Grant number 599175). This research was also supported in part by the Dana-Farber/Harvard Cancer Center Kidney Cancer program, Kidney SPORE P50CA101942-12, and the Trust Family, Michael Brigham, and Loker Pinard Funds for Kidney Cancer Research at Dana-Farber Cancer Institute (all T.K.C.).

## SUPPLEMENTARY NOTE

### Comparison to existing interaction and allele-specific tests

Several alternative strategies exist to leverage allelic imbalance in an interaction framework.

First, the per-sample allelic fraction can be tested for correlation with the tumor/normal indicator across samples. Like any approach that discards information about the individual read count distribution, this suffers from both loss of power and an increased false-positive rate, as shown previously. This bias is particularly severe for low-sample and moderate coverage studies such as ours where the allelic fraction cannot be estimated precisely.

Second, previous studies have used a standard binomial model (equivalent to beta-binomial with no over-dispersion) or a beta-binomial with a single genome-wide over-dispersion parameter (not accounting for local structural variation). To our knowledge, no existing methods model CNA-specific read over-dispersion.

Third, ^21^ proposed a heuristic for calling differential AI when tumor AI is nominally significant (P<0.001) and normal AI is non-significant (P>0.5). While not a formal test, this rule is highly susceptible to false-positives when there is consistent tumor/normal AI but coverage is higher in tumor than normal (leading to greater power and increased significance in tumor). This is especially true for regions of somatically acquired copy number variation, and could lead to the artefactual finding that differential AI is enriched in somatically unstable regions.

Fourth, ^16^ proposed EAGLE, a test for environmental differences in allele-specific expression that accounts for over-dispersion using a generalized linear mixed model. Unlike our method, EAGLE models absolute deviation of allelic imbalance rather than allele-specific deviation. EAGLE is intended for unphased data with heterogeneity of causal effect sizes and was shown to be conservative relative to a directional imbalance model. Due to the substantially different underlying assumptions we did not compare to EAGLE.

Fifth, ^60, 61^ proposed a condition-ASE Z-test based on normal approximations of the ASE parameters. This approximation can be applied to parameters inferred from any ASE model (including stratAS) but is expected to be conservative relative to a formal likelihood test.

Tumor-normal differences can also be evaluated using a traditional eQTL interaction model of the outcome (*y*, for example gene expression) as arising from technical covariates (*c*), baseline expression (β_0_), the genetic variant (*G*), the interaction (*G* * *E*) where *E* is a the tumor/normal indicator, and residual noise: *y* = β_0_ + β_*c*_*c* + β_*g*_*G* + β_*i*_*G*·*E* + ϵ. This model can then be tested for a significant interaction effect β_*i*_≠0. Intuitively, covariates that are strongly correlated with *E* will reduce power to detect a significant interaction by reducing variance between the conditions. In the case where *E* represents tumor/normal data, latent covariates estimated from the expression (such as principal components) generally recapitulate the tumor/normal difference and thus have a severe effect on the power of this test. We do not investigate this test in simulation because its performance relative to AI will be highly dependent on the amount of confounding within/between condition variance. However, multiple previous studies have shown that, even within a single condition, eQTL-based tests are consistently underpowered relative to AI-based tests for sample sizes <100 ^15–17^.

### Figure and Table legends

**Supplementary Figure S1: Power to detect allelic imbalance with stratAS in simulations with 50x upsampled coverage.**

**Supplementary Figure S2: Power and calibration of d-asQTL detection with three methods.** (**a**) QQ plot of asQTL tests for observed coverage (top three panels) and 50x upsampled coverage (bottom three panels). In top panels, the Pearson correlation approach is miscalibrated under the null (AF=0.5) due to poor estimation of the allelic fraction from moderate coverage. (**b**) Power to detect a significant d-asQTL for each method as a function of the tumor allelic fraction. In all simulations the normal allelic fraction is balanced (AF=0.5).

**Supplementary Figure S3: Estimate of cross-state tumor/normal asQTL replication.** (a) red line shows the relationship between discovery FDR and percent of chromatin T-asQTLs that replicate as chromatin nominal N-asQTLs, blue line shows the number of chromatin N-asQTLs that replicate as nominal chromatin T-asQTLs; (b) shows the same relationship for gene expression asQTLs. All quantities estimated using the bootstrapped FDR approach.

**Supplementary Figure S4: Relationship between CNA variability and number of identified asQTLs.** For each state, the genome was ordered from least CNA variable to most and fraction of asQTLs identified (y-axis) is plotted as a function of the fraction of reads (x-axis) for increasing CNA variability. N-asQTL (blue), T-asQTL (red), d-asQTL (green).

**Supplementary Figure S5: # of eGenes identified through asQTL analysis as a function of sample size.** TCGA RNA-seq individuals were downsampled to quantify the number of significant asQTL genes at 5% FDR (blue/green). The same sample subsets were also used to quantify eQTL genes at 5% FDR (orange/red).

**Supplementary Figure S6: Enrichment of per-tissue GTEx eQTLs in asQTL features.** Enrichment for asQTL genes (left) and asQTL peaks (right) for features detected in 5 samples with both RNA-seq and ChIP-seq. Randomly sampled non-asQTL genes/peaks were used as the background.

**Supplementary Figure S7: Estimate of cross-assay tumor/normal asQTL replication.** Top panels show the fraction of gene asQTLs that nominally replicate as chromatin asQTLs; bottom panels show the fraction of chromatin asQTLs that nominally replicate as gene asQTLs. The rows and columns indicate which asQTL state (T-/N-/d-) is considered. All quantities estimated using the bootstrapped FDR approach.

**Supplementary Figure S8: SCARB1 / GRAMD4 response following JQ1 treatment in RCC cell-lines**

**(a)** JQ1 dose response curves of 786-O, A498, and UMRC2 cell lines, showing the different sensitivities of each cell line to JQ1.

**(b)** Measurements of GRAMD4, SCARB1, and MYC RNA transcripts after treatment with JQ1, compared to DMSO controls. *p-value < 0.01, **p-value < 0.001.

**Supplementary Figure S9: Enhancer activity and cell line genotypes at SCARB1 enhancer**

(**a**) Genotyping 786-O RCC cell line at rs4765623 (enhancer) position. Since, 786-O cell line is heterozygous (C/T/T) at rs4765623 this model cell line was chosen for further allelic contribution tests.

(**b**) Genotypes of 786-O triploid cell line at rs5888 (exon8) and rs4765623 (enhancer) position.

(**c**) Measurement of allelic imbalance at rs5888 (exon8) heterozygous reporter position by Sanger sequencing. The allele “A” associates to higher SCARB1 expression.

(**d**) Measurement of allelic imbalance at rs4765623 (enhancer) position. The allele “C” associates higher SCARB1 expression.

(**e**) H3K27ac ChIP was performed and rs4765623 (enhancer) position was amplified from both input and ChIP samples. Amplicons were subjected for Sanger sequencing. According to the altered allelic ratios after ChIP the “C” allele confers higher epigenetic activity.

**Supplementary Figure S10: Heritability enrichment for intersections of functional features.**

**Supplementary Table S1: Correlation of local overdispersion parameter and CNA fold change for each analyzed sample in tumor and normal.**

**Supplementary Table S2: Coefficients from logistic regression model of balance/imbalance as a function of local features.**

**Supplementary Table S3: Replication rate of chromatin and gene expression asQTLs identified in individual samples.**

**Supplementary Table S4: Super-enhancers identified at lead RCC GWAS SNPs.**

**Supplementary Table S5: Gene expression asQTLs identified at lead RCC GWAS SNPs.**

**Supplementary Table S6: Joint eQTL & GWAS fine-mapped SNPs in the SCARB1 locus.**

**Supplementary Table S7: RCC GWAS heritability enrichment for analyzed functional features.**

**Supplementary Table S8: Mean GWAS heritability enrichment from 95 public traits for analyzed functional features.**

**Supplementary Table S9: Enrichment of GWAS heritability for each of 95 public traits and each analyzed feature.**

**Supplementary Table S10: Significance of enrichment of GWAS heritability for each of 95 public traits and each analyzed feature.**

**Supplementary Table S11: Enrichment of six RCC-relevant TFs in H3k27ac peaks.**

**Supplementary Table S12: Enrichment of six RCC-relevant TFs in asQTL peaks relative to matched peaks (top) and all peaks (bottom).**

**Supplementary Table S13: Enrichment of six RCC-relevant TFs in T-/d- asQTL peaks relative to exclusively N-asQTL peaks.**

**Supplementary Table S14: Overlap of HIF binding elements at RCC GWAS lead SNPs. Supplementary Table S15: Oligonucleotide, gRNA and shRNA sequences.**

