## Supplementary figures and images for "Allelic imbalance reveals widespread germline-somatic regulatory differences and prioritizes risk loci in Renal Cell Carcinoma"

### Supplemental Figures

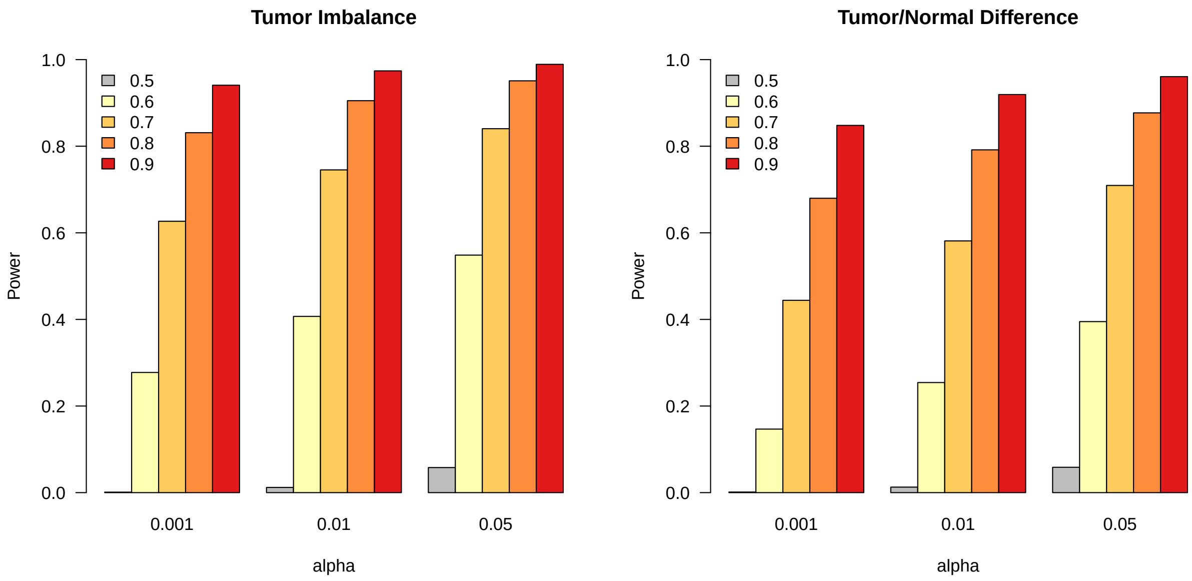

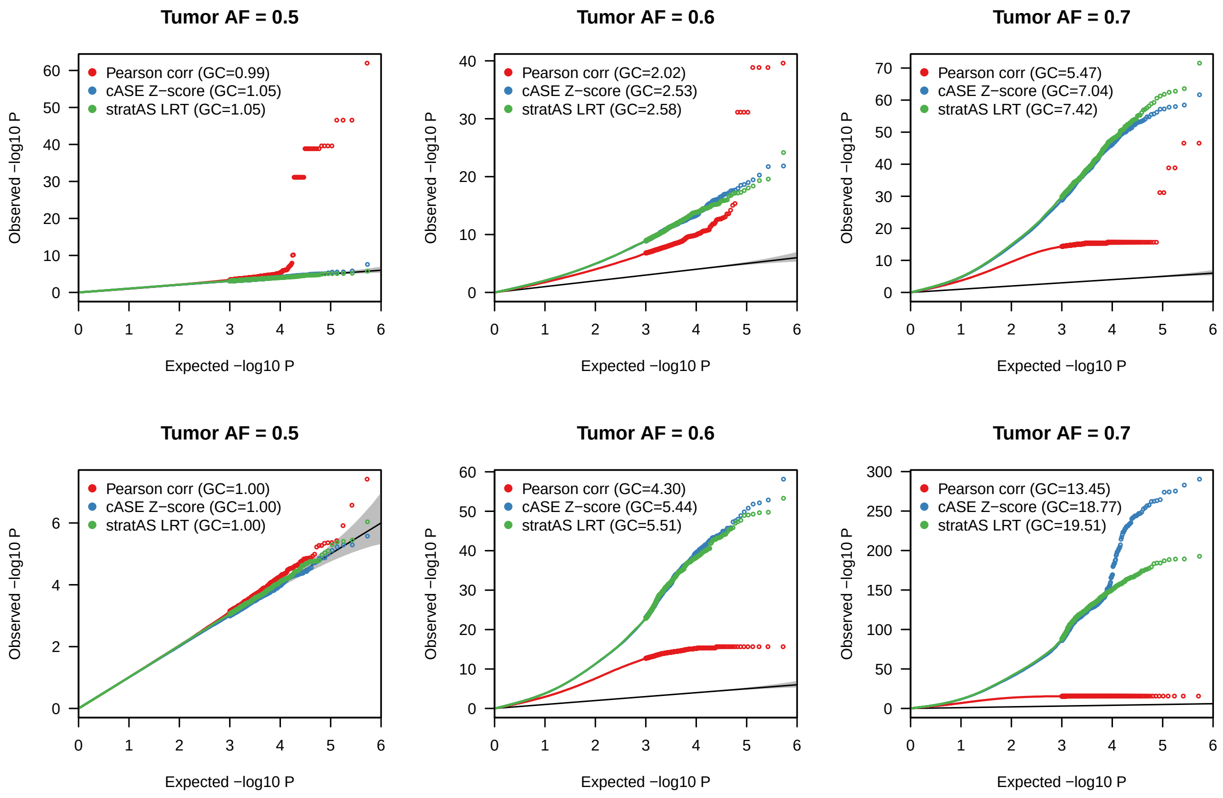

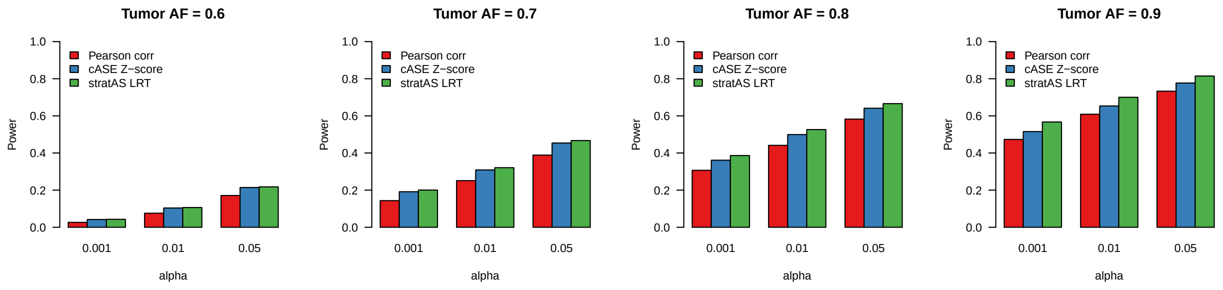

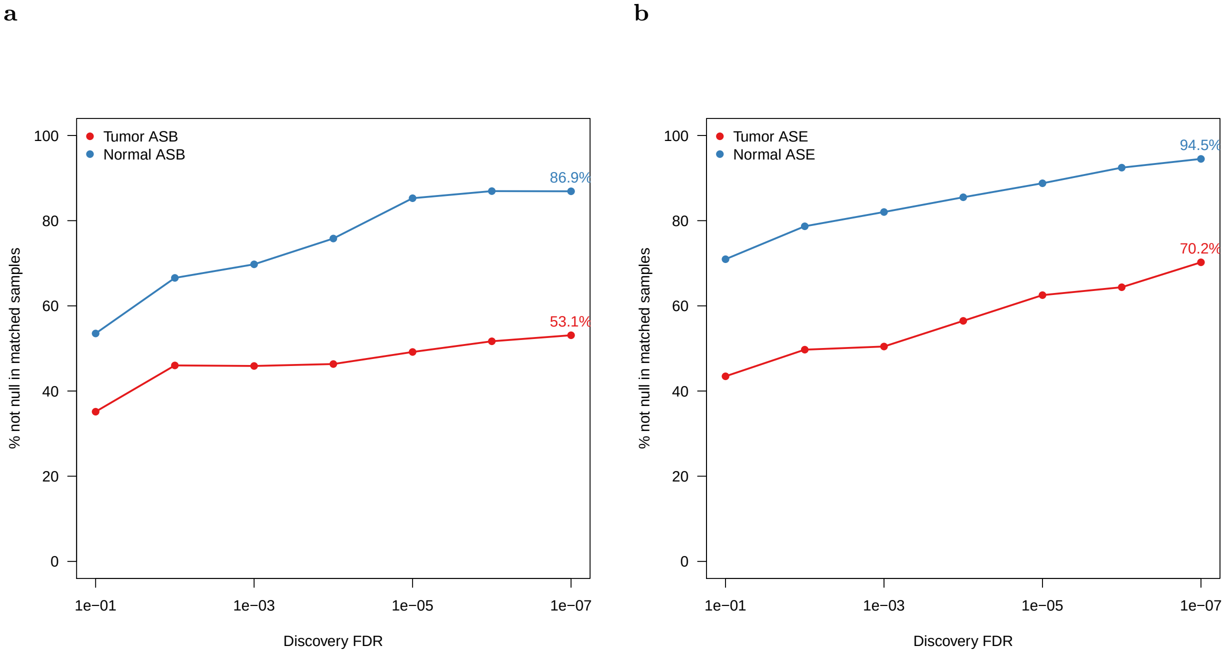

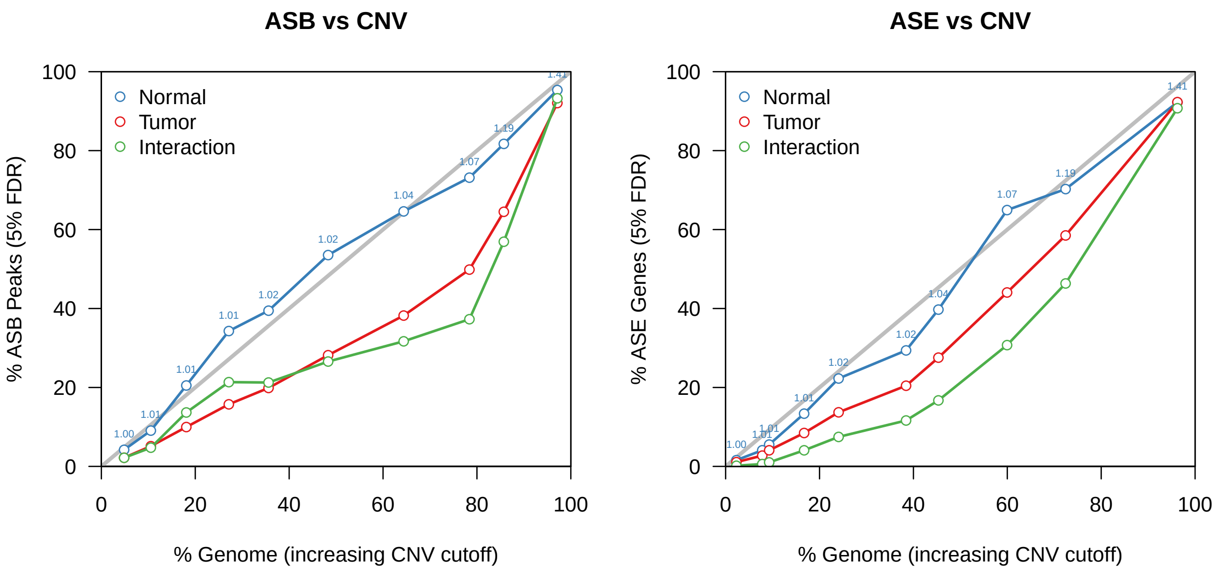

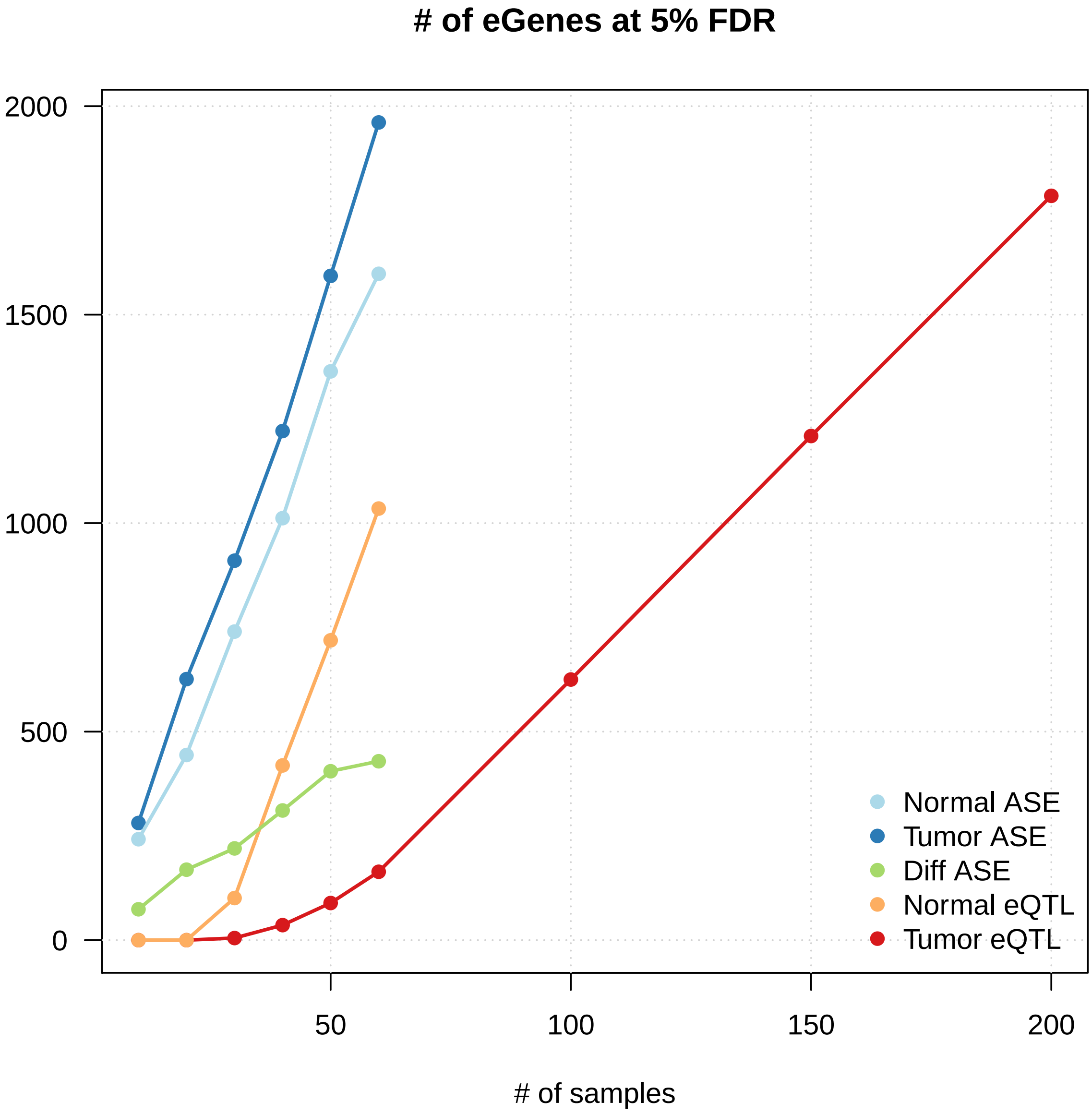

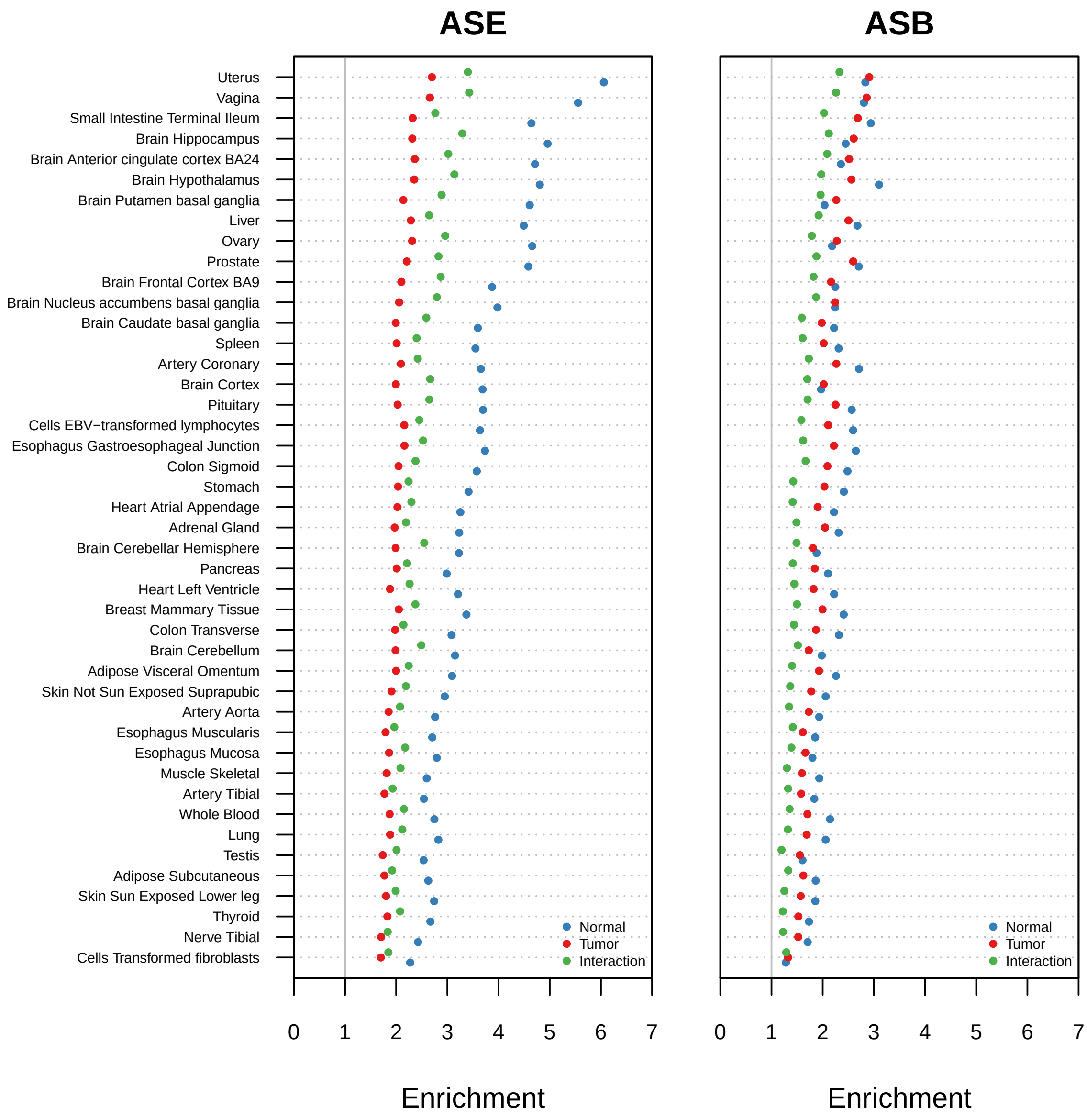

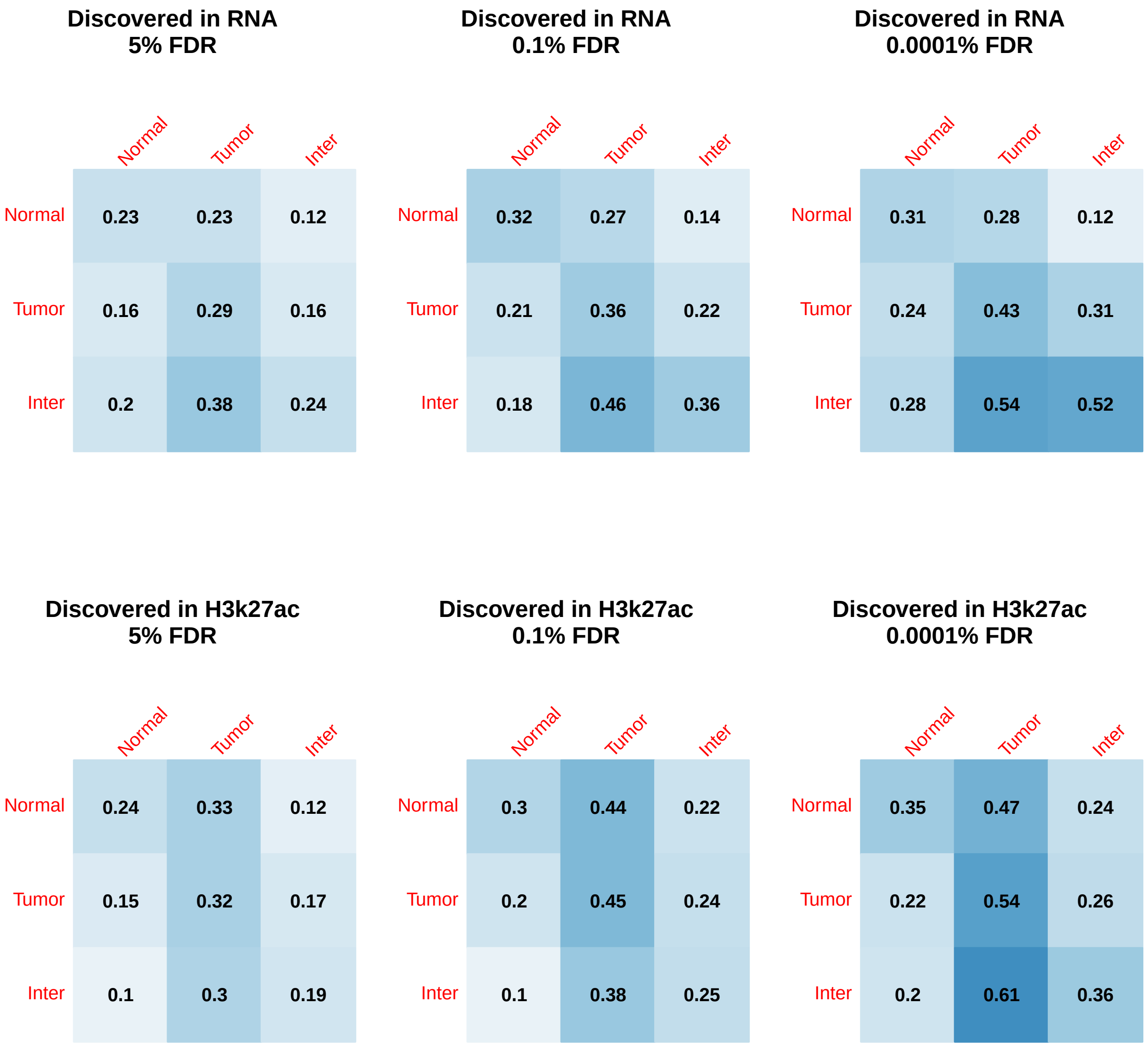

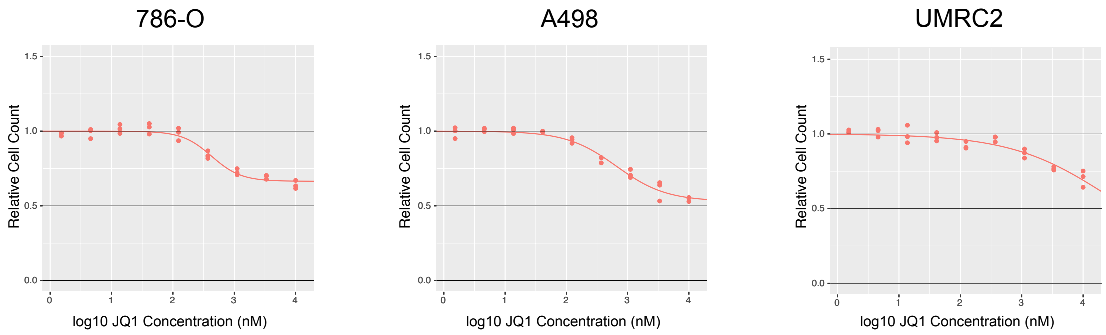

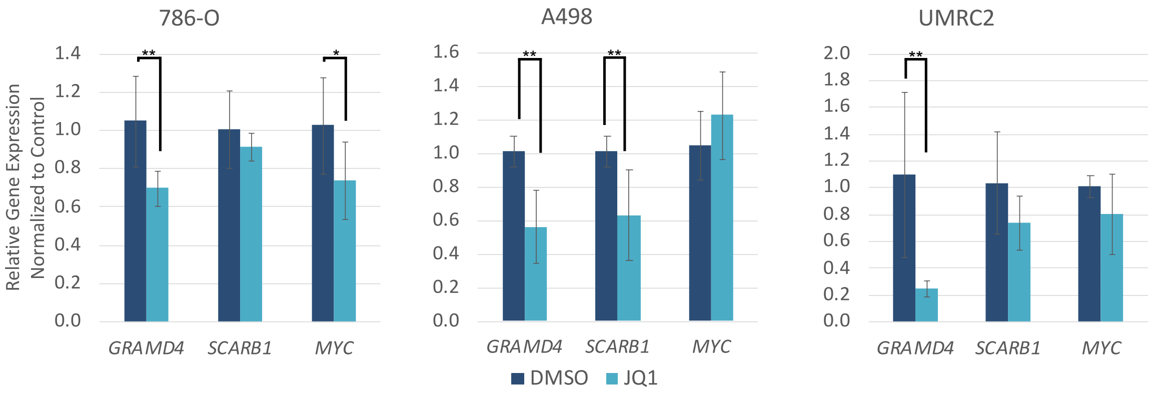

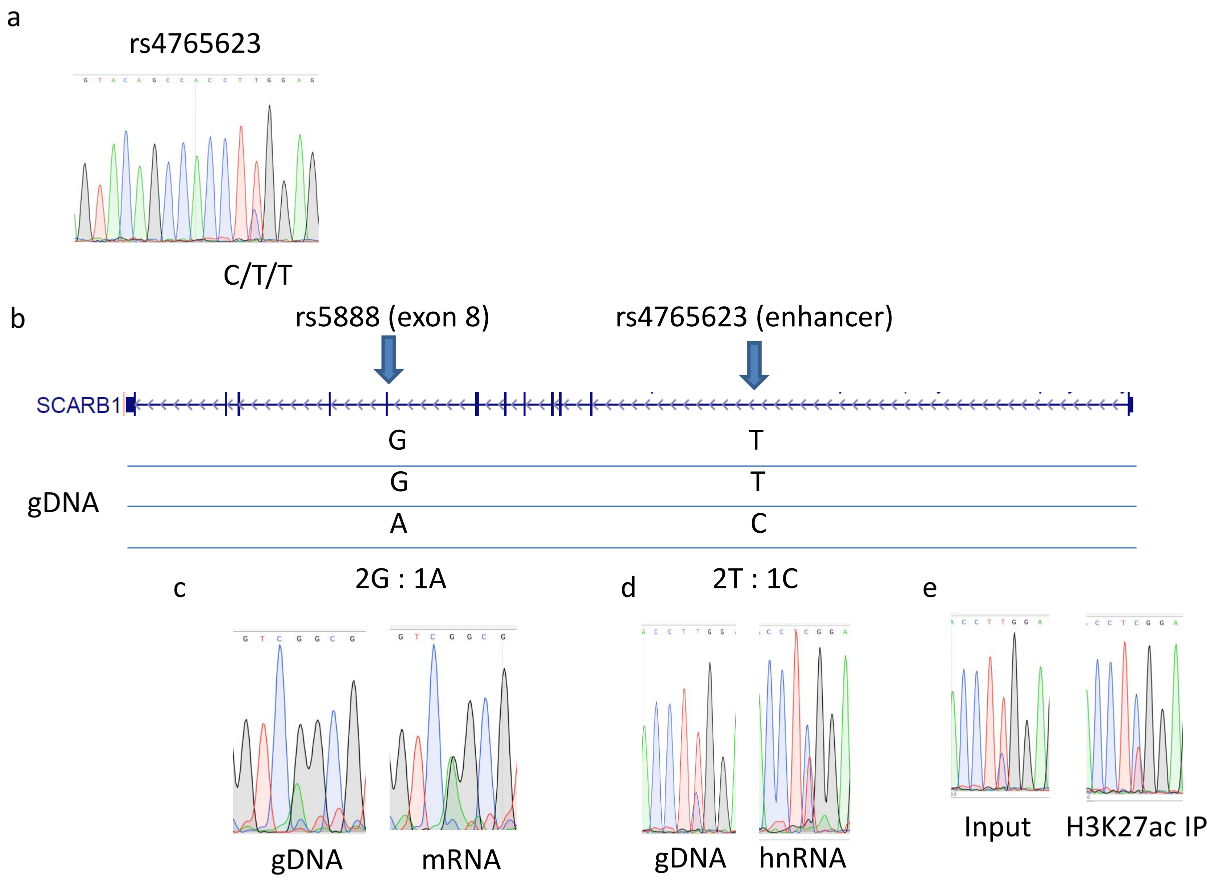

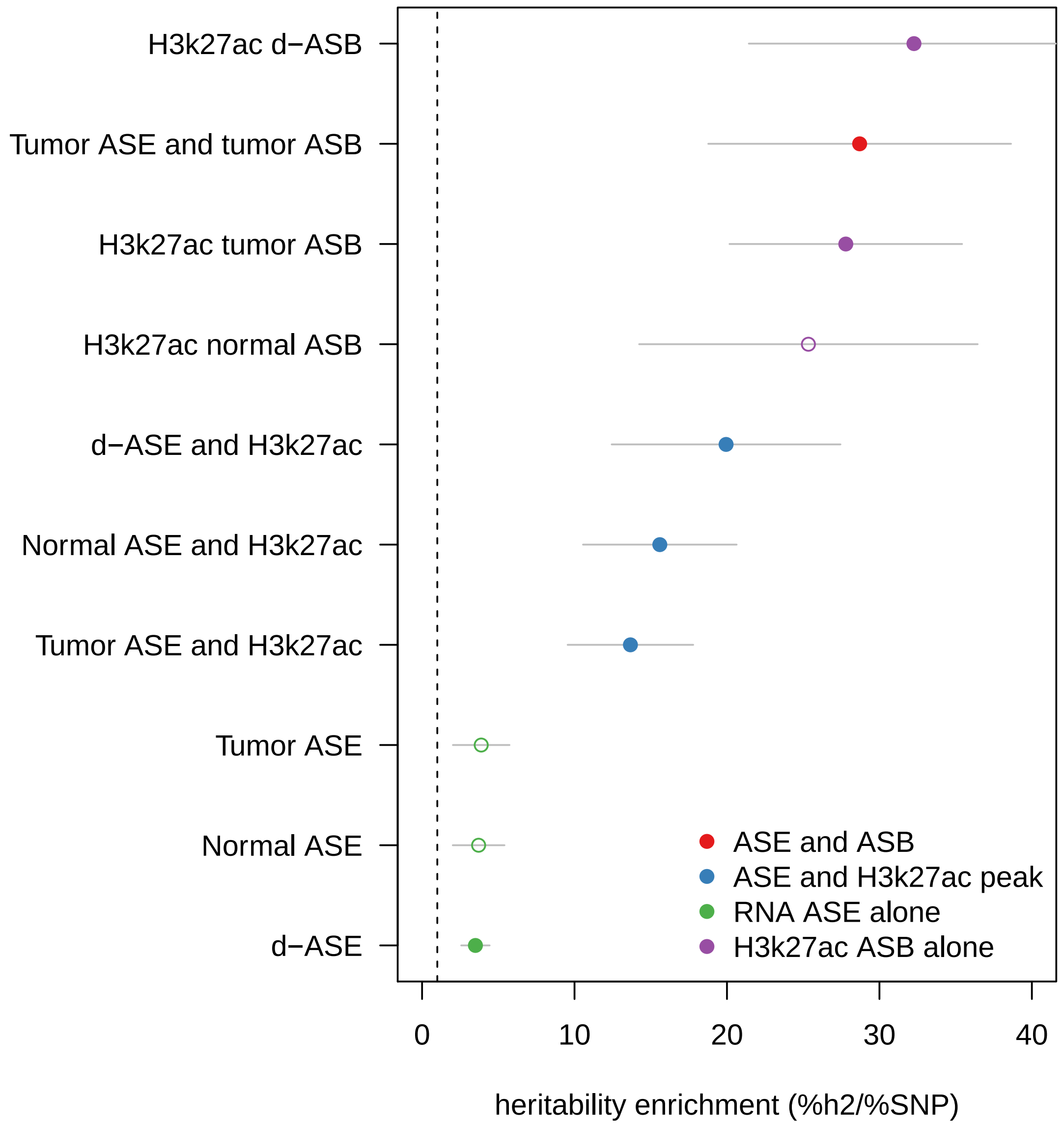
